# Pervasive ethanol in the fruits and floral nectars consumed by tropical frugivores

**DOI:** 10.64898/2026.11.11.766018

**Authors:** Julia G. Casorso, Konstans Wells, Allegra N. DePasquale, Anna C. Bowland, William D. Aguado, Sylvia Alsisto, Erika B. Alves, Marc Ancrenaz, Francisca S. Andicene, Maria Anjum, Filippo Aureli, Madelyn A. Barnes, Elena Bersacola, Monika Bisht, Susana Carvalho, Adriano A. B. Darosci, Terita R. Deare, Anthony Di Fiore, Nathaniel J. Dominy, Gideon A. Erkenswick, Elodie Freymann, Taniya Gill, Benoit Goossens, Luja Guju, Goro Hanya, Xavier A. Harrison, Laura T. Hernández Salazar, Nadine Hussein, Steig E. Johnson, Fumiya Kakuta, Rentaro Kimpara, Yosuke Kurihara, Wanyi Lee, Shelly Masi, Miku Minamikawa, Lara Nellissen, Omer Nevo, Julia Ostner, Melody A. Petersen-Leong, Alice C. Poirier, Ishika Ramakrishna, Hasinala Ramangason, Carlos E. Ramírez Torres, Diary N. Razafimandimby, Leticia V. Rodrigues, Suheidy Romero Morales, Jessica M. Rothman, Oliver Schülke, Fred Y. Y. Tuh, Atsushi Usami, Sri Suci Utami Atmoko, Vivek V. Venkataraman, Erin R. Vogel, Mrinalini Watsa, Matthew A. Carrigan, Kimberley J. Hockings, Amanda D. Melin

**Author notes:** Bernhard Nocht Institute for Tropical Medicine (BNITM), Ethnopharmacology and Zoopharmacognosy, Hamburg, Hamburg, Germany (Elodie Freymann). Ashoka Trust for Research in Ecology and the Environment (ATREE), Bengaluru, Karnataka, India (Taniya Gill). Institute of Life and Environmental Sciences, University of Tsukuba, Tsukuba, Ibaraki, Japan (Wanyi Lee). Department for the Ecology of Animal Societies, Max Planck Institute of Animal Behavior, Konstanz, Germany (Lara Nellissen). Alberta Biodiversity Monitoring Institute, University of Alberta, Edmonton, Alberta, Canada (Melody A. Petersen-Leong). Université Bourgogne Europe, Institut Agro, CNRS, INRAE, UMR CSGA, Development of Olfactory Communication & Cognition Laboratory, Dijon, France.

## Abstract

Dietary ethanol has long been proposed as a selective force shaping the diets and physiology of tropical frugivores and nectarivores, yet this idea has rested on measurements from only a few species and sites. Here, we present the first global survey of ethanol in wild fruits and flowers (591 and 37 species, respectively, across 18 countries) and test ecological and phylogenetic predictors of its variation. We detected ethanol in the ripened fruits of 72% of angiosperm species (211 species, *N* ≥ 3 per species; 0.021% - 3.9% alcohol by volume (ABV)). Ethanol was also detected in the flowers of 12 species (0.037% - 0.73% ABV). Maximum fruit ethanol varied with temperature, humidity, and phylogenetic relatedness, and it was higher in ripe fruits collected from the forest floor relative to their canopy counterparts. We then extrapolated potential daily ethanol consumption for 10 primate species. Estimated maximum daily ethanol ingestion for primates ranged from negligible to 0.72 g/kg/day on average (equivalent to 5.1 international standard drinks, 10 g ethanol each, for a 70 kg human). Our findings show that ethanol is widespread in plant reproductive tissues and shaped by both phylogeny and environment, supporting the hypothesis that dietary ethanol has been a pervasive selective pressure on fruit- and nectar-feeding animals.

## INTRODUCTION

Natural ethanol (alcohol) is an overlooked component of plant–animal interactions that may influence the ecology, behaviour, and evolution of fruit- and nectar-feeding animals (*1–3*). Ethanol is produced when naturally-occurring yeasts ferment sugars, a process that varies with fruit and nectar chemical composition, environmental conditions, and microbial communities (*1*, *4–6*). Because ethanol can provide an energetic resource while also producing physiological and behavioural effects that can impact health, its availability in plant reproductive tissues may represent an overlooked ecological dimension of animal diets (reviewed in (*1*)). Yet, despite decades of interest and debate regarding the presence and evolutionary significance of natural ethanol, empirical knowledge remains largely restricted to small-scale studies of a limited number of plant species (*1*, *2*, *7*, *8*). Consequently, the extent to which ethanol is a widespread feature of plant reproductive tissues, and the ecological and evolutionary processes shaping its distribution, remain largely unknown.

The potential importance of dietary ethanol arises from its dual role as both a resource and a physiological challenge. At low to moderate concentrations, ethanol consumption may provide readily metabolizable energy, stimulate appetite, and influence feeding decisions, whereas these potential benefits are countered by excessive consumption that can result in intoxication, disorientation, and other toxic effects (*9–11*). Accordingly, variation in ethanol availability may impose varied selective pressures on fruit- and nectar-feeding animals, favouring adaptations that enable sensory detection, tolerance, metabolism, or avoidance of ethanol-rich foods (*12–15*).

Non-human primates (hereafter primates) have featured prominently in discussions of dietary ethanol as many species are highly frugivorous, and their response to ethanol may provide insights into the evolutionary origins of human alcohol use (*2*, *16–18*). For example, a single amino acid substitution in the alcohol dehydrogenase class IV gene (*ADH4*) in the last common ancestor of African apes increased the enzyme’s catalytic efficiency for ethanol metabolism approximately forty-fold (*16*). This ancient adaptation is hypothesised to have facilitated increased human consumption of fermented food and drink during the Neolithic transition (*17*). Scrumping of fruits from the forest floor, which may contain more ethanol than canopy fruits, has featured repeatedly in dialogue on *ADH4* evolution but not been tested directly (*16*, *19*). Furthermore, ethanol may influence contemporary primate foraging behaviour: individuals frequently assess fruit quality through olfactory cues (*20–22*), show high sensitivity to ethanol-associated odours (*14*), and some species (slow loris, *Nycticebus coucang*; aye-aye, *Daubentonia madagascariensis*) tolerate or even favour moderate ethanol concentrations in experimental settings (*23*). Recent field studies have detected ethanol at concentrations likely to have biological impact in fruits consumed by spider monkeys (*Ateles geoffroyi*) and chimpanzees (*Pan troglodytes*) (*24*, *25*), and ethanol may contribute to social aspects of feeding among wild chimpanzees (*26*). However, whether observations of wild animals ingesting ethanol represent isolated examples or a widespread feature of tropical and subtropical plant–animal interactions remains unresolved.

Ethanol availability should vary predictably across environments and plant lineages: temperature and humidity influence yeast growth and fermentation (*4*, *27*, *28*), and phylogenetically preserved fruit traits (e.g., sugar content, mesocarp structure) may cause ethanol concentration to covary with plant relatedness (*29*, *30*). However, existing evidence remains too taxonomically and geographically fragmented to establish the broad-scale drivers of ethanol variation.

Here, we quantify ethanol in 4,173 fruits from 591 angiosperm species found in natural forests and agricultural mosaics across 18 countries and six major biogeographical realms (Afrotropical, Madagascan, Neotropical, Oriental, Panamanian, and Sino-Japanese (*31*); Fig. 1, supplementary text S1). We then focus on a subset of species for which the ripened (i.e., excluding unripe) fruit were able to be sampled repeatedly (*N* ≥ 3), comprising 2,575 fruit samples from 211 species across 29 sampling locations (supplementary text S2, fig. S1). We also measured floral tissues of 39 wild angiosperms. We pursued three main objectives: (1) to quantify the prevalence of ethanol across plant reproductive tissues and assess variation within and among species; (2) to test whether environmental variables and phylogenetic relationships among plant species predict ethanol occurrence and concentration; and (3) to determine whether fruits occurring in the canopy compared to fruit collected from the forest floor differ in ethanol concentration. We predicted that ethanol would occur broadly across plant reproductive tissues and reach concentrations likely to influence animal physiology and behaviour; that ethanol occurrence and concentration would increase under warmer and more humid conditions that favour microbial fermentation; and that fallen fruits would contain higher ethanol concentrations than canopy fruits because fruits reaching the forest floor are typically riper and experience greater exposure to conditions promoting fermentation.

**Fig. 1.**
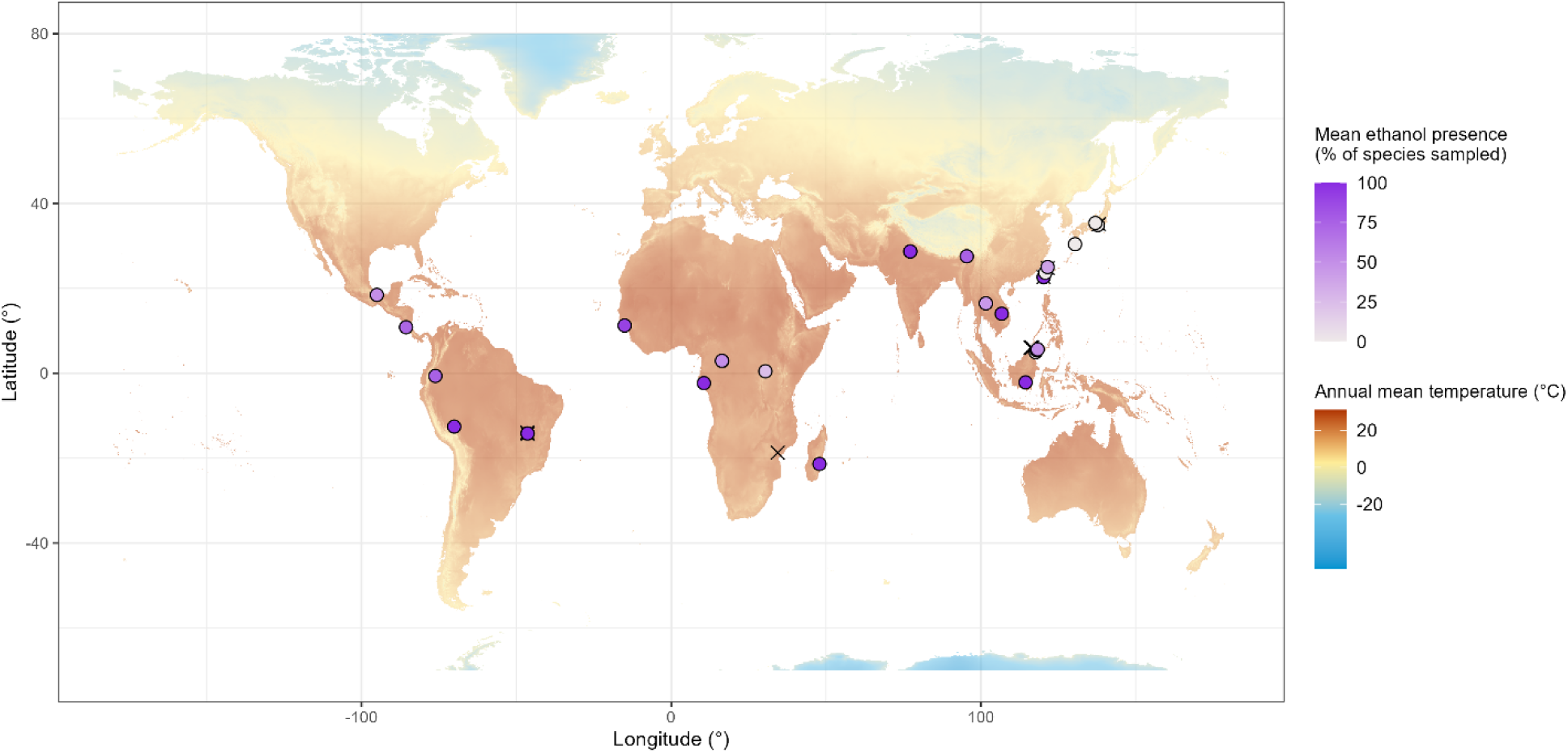
Global map depicting all sampling sites (*N* = 39) where fruits were sampled for this study. For sites with sufficient sampling repetition (i.e., at least three samples per species, excluding unripe fruits; *N* = 29), the colour gradient of points represents the mean ethanol presence (% of species sampled) for different fruit species at each site, ranging from ethanol detected in 0% (light grey points) to 100% (dark purple points) of species sampled. Sites depicted by an X (*N* = 10) were excluded from the repeatedly sampled dataset used for statistical analyses due to inadequate sampling repetition. The background color gradient on the map depicts annual mean temperature, ranging from blue for low temperatures to red for high temperatures (obtained from the WorldClim database version 2.1 (*59*)).

Finally, to place plant-level ethanol measurements into the context of potential animal dietary exposure, we estimated maximum daily dietary ethanol intake for 10 wild primate species as an exemplar group of tropical frugivores. We focus on primates because they are the most prevalent nonvolant frugivorous taxon in the tropics and have a long evolutionary history with angiosperms (*32–34*). By combining global measurements of plant ethanol availability with phylogenetic, environmental, and dietary analyses, we provide the first broad-scale framework for understanding natural ethanol exposure and evaluate its potential role as an ecological and evolutionary force shaping interactions between plants and fruit- and nectar-feeding animals.

## RESULTS

### Ethanol occurrence in wild fruits and flowers

In the full dataset of sampled fruits, which included all ripeness stages across fruit of all sampled angiosperm species (*N* = 591 species; fig. S1), ethanol was detected in 272 of the 591 species (46.0%). However, excluding unripe fruits, ethanol was widespread and detected in 71.6% of the 211 fruit species that were repeatedly sampled (table S1). This difference reflects the lower sampling effort in some species, and the inclusion of unripe fruits, in the full dataset. At the individual fruit level in the repeatedly sampled dataset, ethanol was detected in 69.7% of overripe fruits (*N* = 409 fruits), 47.0% of ripe fruits (*N* = 1,744), and 33.6% of midripe fruits (*N* = 438). Among non-zero measurements, mean ethanol concentration was 0.27% alcohol by volume (ABV). Maximum concentrations were 3.90% ABV in overripe fruits, 3.75% ABV in ripe fruits, and 3.00% ABV in midripe fruits. The maximum concentration for unripe fruits (i.e., from the full fruit dataset) was 1.49% ABV (fig. S2). Overall, the majority (i.e., central 90%) of all non-zero values in the full fruit dataset fell between 0.03 and 1.05% ABV.

Ethanol-containing fruits occurred across a wide diversity of angiosperm lineages, including widespread tropical plant families such as Arecaceae, Fabaceae, Moraceae, Myrtaceae, and Rubiaceae (table S1). The ten fruit species with the highest recorded ethanol concentrations in our dataset are presented in Table 1, with several additional taxa exceeding 1% ABV (table S1).

**Table 1.** The top 10 fruit species with the highest recorded ethanol concentrations in our study. Maximum ethanol values (EtOH_max_; % alcohol by volume (ABV)) represent the highest reliable ethanol concentration that was measured across all sampled fruits for each species. Each species is listed along with their taxonomic family and the ripeness stage from which the sample with the highest measurement was obtained. Note the *Anacardium occidentale* sample was a wadge of multiple *A. occidentale* fruits made by a chimpanzee.

| Rank | Species | Family | EtOH<sub>max</sub><br>(% ABV) | Sample ripeness stage |
|---|---|---|---|---|
| 1 | *Spondias mombin* | Anacardiaceae | 3.90 | Overripe |
| 2 | *Anacardium occidentale* | Anacardiaceae | 3.75 | Ripe |
| 3 | *Mangifera indica* | Anacardiaceae | 3.22 | Overripe |
| 4 | *Treculia africana* | Moraceae | 2.96 | Ripe |
| 5 | *Spondias radlkoferi* | Anacardiaceae | 2.92 | Ripe |
| 6 | *Salacia crassifolia* | Celastraceae | 2.86 | Ripe |
| 7 | *Borassus aethiopum* | Arecaceae | 2.01 | Ripe |
| 8 | *Solanum hazenii* | Solanaceae | 1.99 | Ripe |
| 9 | *Parinari anamensis* | Chrysobalanaceae | 1.96 | Ripe |
| 10 | *Musa* sp. | Musaceae | 1.91 | Overripe |

Floral tissues were sampled less extensively and were analysed descriptively. Ethanol was detected in 12 of 37 species where we sampled flowers or flower-buds (32%), with non-zero concentrations ranging from 0.04 to 0.37% ABV (table S2). The fruits of three of these 12 species were also represented in the repeatedly sampled dataset, and all had measurable fruit ethanol. We additionally sampled 12 floral nectar samples and 14 samples of liquid accumulated within the nectar-containing bracts of the traveller’s tree (*Ravenala madagascariensis*) in Madagascar. Ethanol concentrations reached up to 0.73% ABV in floral nectar and 0.43% ABV in accumulated bract liquid (fig. S3)

### Phylogenetic signal

Maximum ethanol concentration (EtOH_max_) in the repeatedly sampled dataset (*N* = 211; fig. S1) showed a phylogenetic signal (Fig. 2). Pagel’s λ for EtOH_max_ was 0.697 and differed significantly from both λ = 1 (*P* < 2.2 × 10□¹□) and λ = 0 (*P* = 5.4 × 10□□), consistent with an intermediate phylogenetic signal. If we examine the full set of taxonomically identified fruit species (*N* = 583; fig. S1, table S1), we similarly detected moderate phylogenetic signal in EtOH_max_ (Pagel’s λ = 0.496; significantly different from λ = 1, *P* < 2.2 × 10□¹□, and λ = 0, *P* = 6.7 × 10□¹□; fig. S4). In contrast, we found no evidence of a phylogenetic signal in mean (EtOH_mean_) or median (EtOH_median_) ethanol concentrations, with Pagel’s λ estimates equal to zero in both analyses.

**Fig. 2.**
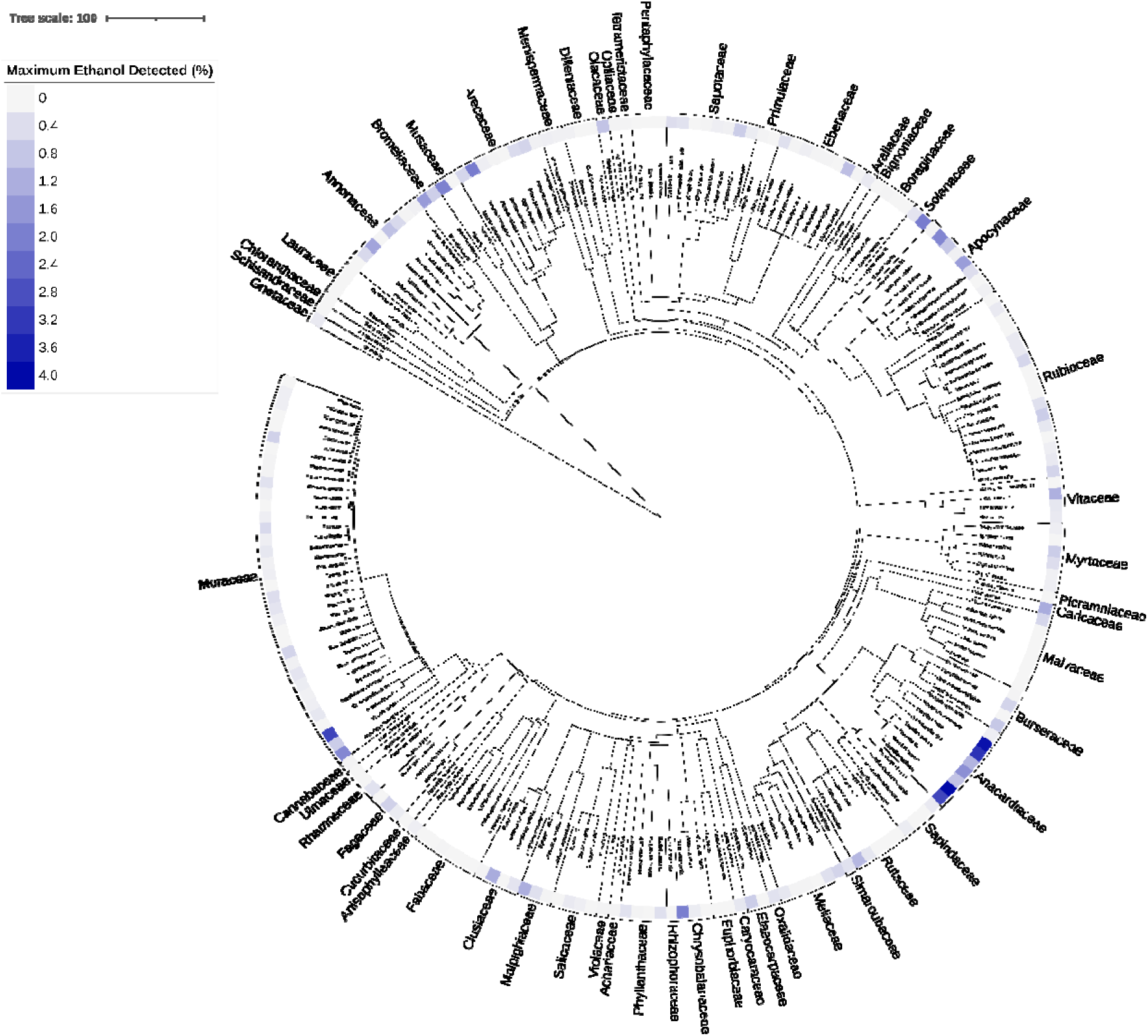
Phylogenetic tree of fruit species and their highest ethanol concentrations detected. This tree includes all 211 species (classified at least to taxonomic genus) in the repeatedly sampled dataset (i.e., at least three samples per species, excluding unripe fruits). The maximum ethanol concentration (% alcohol by volume) detected for each species is visualized along a colour gradient, with darker colours representing higher values. Brackets indicate taxonomic family. The scale reflects a distance of 100 Ma. This tree was generated using the mega-tree ‘GBOTB.extended.WP.tre’ in V.PhyloMaker2 (*56*) in R version 4.6.0 (*62*), and visualized with Interactive Tree of Life (iTOL) version 6 (*63*).

### Environmental predictors of fruit ethanol occurrence and concentration

Fruit ethanol occurrence varied nonlinearly with environmental conditions (Fig. 3; table S3). Ethanol occurrence (EtOH_occurrence_) increased with annual mean temperature up to 25°C, reaching a predicted peak occurrence frequency of 83.2% (95% confidence interval (CI): 70.0–91.4%), before declining at higher temperatures (Fig. 3A). EtOH_occurrence_ further showed nonlinear, saturating relationships with both mean annual precipitation and precipitation seasonality, peaking at 3,335 mm annual precipitation (estimated EtOH_occurrence_ = 95.3%, 95% CI: 74.1–99.3%; Fig. 3B) and at a precipitation seasonality coefficient of variation of 133.1% (estimated EtOH_occurrence_ = 88.1%, 95% CI: 49.0– 98.3%; Fig. 3C). EtOH_occurrence_ also increased linearly with vegetation density (i.e., normalized difference vegetation index, NDVI) until NDVI values of 0.4, remained relatively constant between NDVI values of 0.4 and 0.6, and declined somewhat at higher values. Predicted occurrence was highest at an NDVI of approximately 0.5 (estimated EtOH_occurrence_ = 85.2%, 95% CI: 65.6–94.6%; Fig. 3D).

**Fig. 3.**
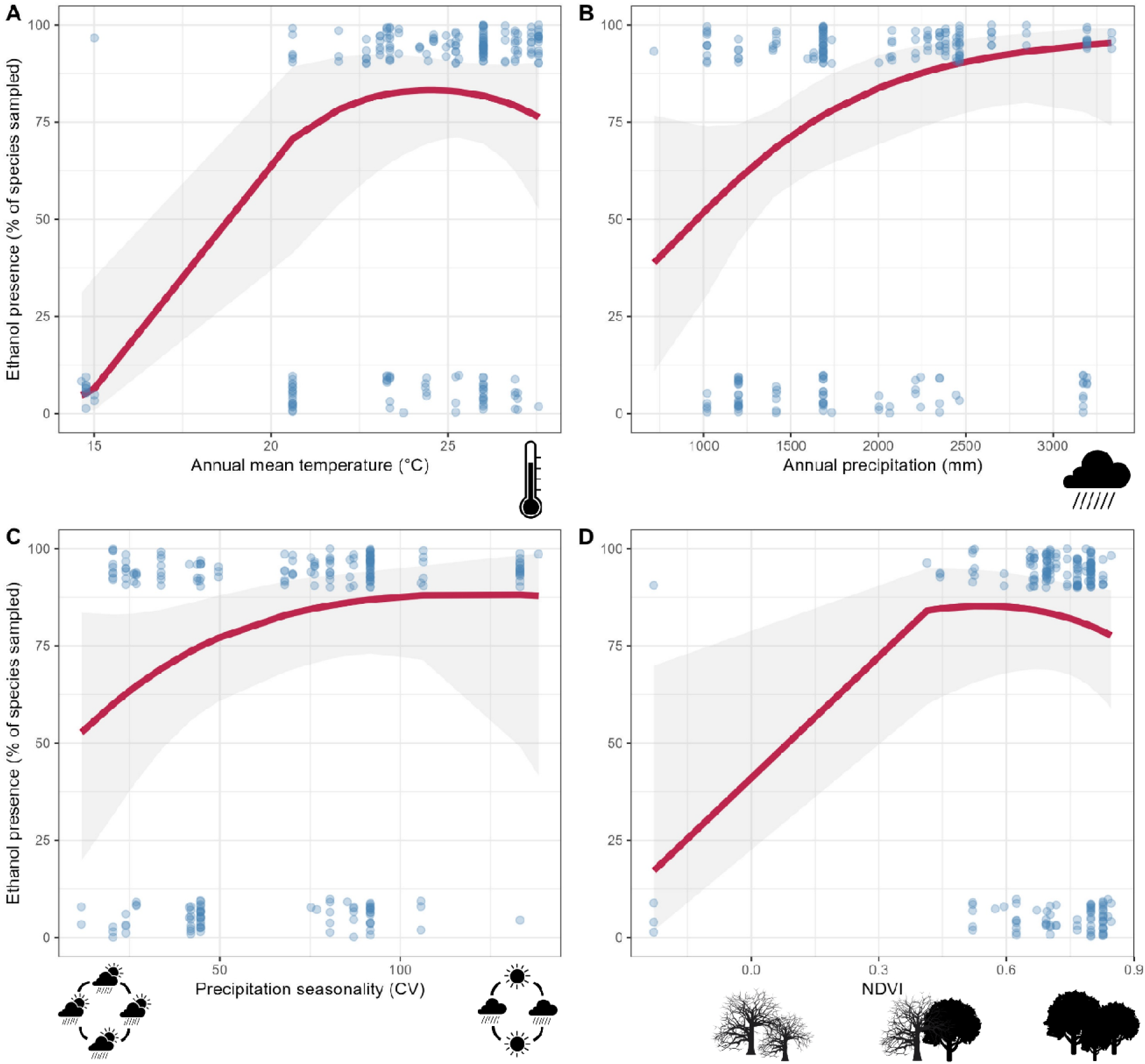
Plots of predicted relationships between fruit ethanol occurrence and significant environmental variables. Each plot depicts fruit ethanol presence (% of species sampled that had ethanol detected in at least one reliable measure across our entire raw dataset) as a function of **(A)** annual mean temperature (°C), **(B)** annual precipitation (mm), **(C)** precipitation seasonality (coefficient of variation, CV), and **(D)** normalized difference vegetation index (NDVI) across sites, for the repeatedly sampled dataset (i.e., at least three samples per species, excluding unripe fruits, *N* = 211 species). Curves and shaded areas represent the predictions and their pointwise 95% confidence intervals, respectively, of the most supported model (selected based on the corrected Akaike information criterion (AIC_c_)).

Points represent the raw data associated with the model’s binary dependent variable of ethanol presence (true/false, shown as 100% or 0% on the plot, respectively) and the independent variable shown on each plot, with an introduced vertical error of magnitude 5 units to improve visualization (darker colours represent greater overlap of points). Annual mean temperature, annual precipitation, and precipitation seasonality data were obtained from the WorldClim database (version 2.1 (*59*)) and NDVI data from MODIS satellite data (MODIS MOD13Q1 version 6 (*60*)).

Environmental predictors of ethanol concentration differed partially from those of ethanol occurrence (Fig. 4, tables S4–S5). Both EtOH_mean_ and EtOH_max_ showed nonlinear relationships with mean annual precipitation, reaching predicted maxima at 3,335 mm. At this value, estimated EtOH_mean_ was 0.46% ABV (95% CI: 0–0.97%) and estimated EtOH_max_ was 1.39% ABV (95% CI: 0–2.83%; Fig. 4A). EtOH_mean_ and EtOH_max_ also showed nonlinear relationships with precipitation seasonality, reaching predicted maxima at a seasonality coefficient of variation of 91.7%. At this value, estimated EtOH_mean_ was 0.17% ABV (95% CI: 0.12–0.23%) and estimated EtOH_max_ was 0.54% ABV (95% CI: 0.36–0.71%; Fig. 4B). Unlike EtOH_occurrence_, annual mean temperature and vegetation density were not significantly associated with EtOH_mean_ or EtOH_max_.

**Fig. 4.**
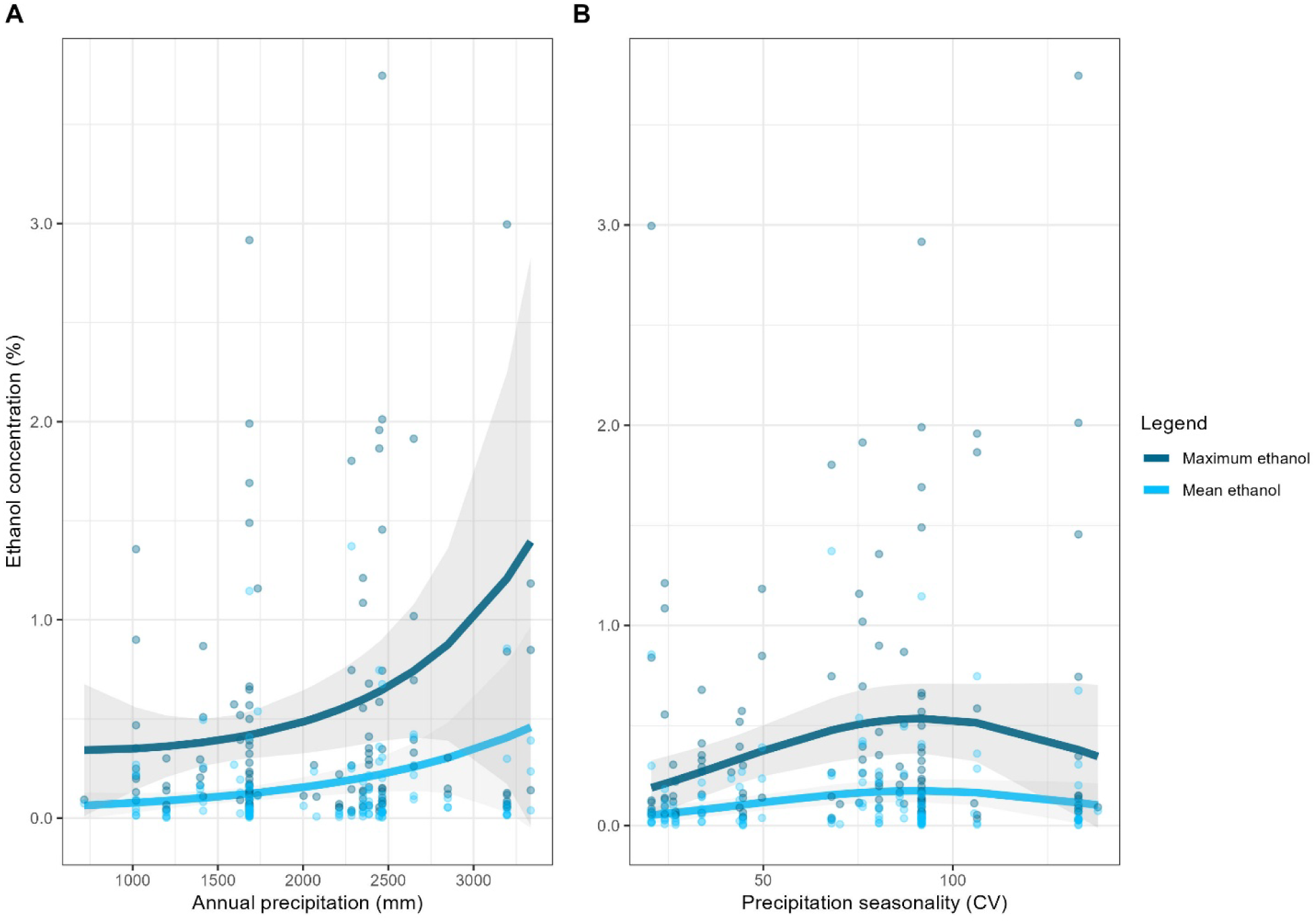
Plots of predicted relationships between fruit ethanol concentration and significant environmental variables. Each plot depicts the mean and maximum fruit ethanol concentration (% alcohol by volume) as a function of **(A)** annual precipitation (mm) and **(B)** precipitation seasonality (coefficient of variation, CV) across sites, for the repeatedly sampled dataset (i.e., at least three samples per species, excluding unripe fruits, *N* = 211 species). Curves and shaded areas represent the predictions and their pointwise 95% confidence intervals, respectively, of the two models with the greatest support across both mean and maximum ethanol concentration (selected based on the corrected Akaike information criterion (AIC_c_)). Points represent the raw data used in the models. Environmental data were obtained from the WorldClim database (version 2.1 (*59*)).

### Sampling fruits from the canopy versus ground

To estimate whether fruits collected from the forest floor had higher ethanol content than those in the canopy, we compared ethanol concentrations of ripened fruits of 68 plant species where fruits were sampled from both the ground and canopy. Ground-collected fruits had significantly greater ethanol content compared to canopy-collected fruits (one-tailed *Z* test, EMM ± SE = 0.039 ± 0.020%, *Z* ratio = 195.9, *P* = 0.025; Fig. 5). We further tested if there was an effect of ground versus canopy collection site by ripeness stage by running a separate model including fruit ripeness category. We found a significant interaction between ripeness stage and collection site (Type III Wald chi-square test, χ^2^(2) = 14.08, *P* = 8.7 x 10^-4^; table S6). Pairwise comparisons (with Holm-Bonferroni correction) indicated that ripe fruits collected from the ground had 44% higher odds of having greater ethanol concentrations than those collected from the canopy (two-sided Wald *Z* test, odds ratio = 1.44, corrected *P* < 1.0 x 10^-4^, 95% CI: 1.19–1.74). For midripe and overripe fruits, there was no significant difference in the odds of canopy and ground collected fruits differing in ethanol concentration (corrected *P* = 0.38 for both; table S7).

**Fig. 5.**
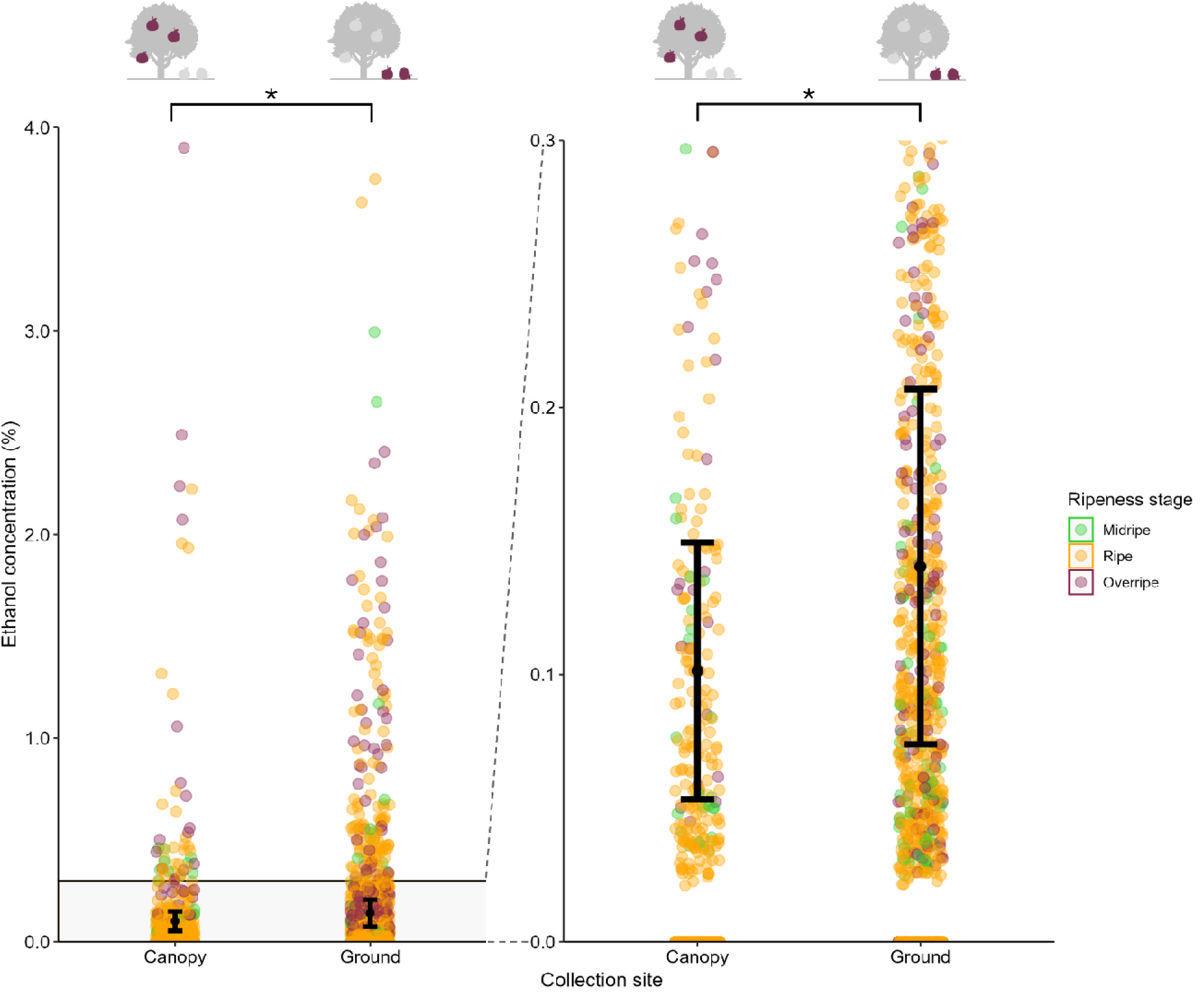
Comparison of ethanol concentration in fruits collected from the canopy versus the forest floor. The lefthand panel depicts the entire distribution of ethanol concentration (% alcohol by volume (ABV)) of ripened fruits from 68 species sampled from both the canopy (N = 455 fruits) and ground (N = 676 fruits). An enlarged view of the greyed section (between 0 to 0.3% ABV) is shown in the righthand panel. Data are from the repeatedly sampled dataset (i.e., at least three samples per species, excluding unripe fruits), further filtered for species with (a) ethanol detected in at least one reliable measure, and (b) at least one fruit sampled from the ground and the canopy. Data points represent individual sample measurements, coloured according to ripeness stage (midripe, ripe, or overripe), with an introduced error of 0.1% to improve visualization of overlapping points. Black bars depict estimated marginal means (EMMs, ± standard error) on the response scale, with an asterisk denoting statistical significance (α = 0.05). EMMs were derived from the model investigating the impact of collection site on ethanol concentration. Note the gap of missing points between 0 and 0.02% ABV is a consequence of the breathalyzer’s minimum detection limit.

### Predicted dietary ethanol exposure in primates

To estimate potential dietary impacts of fruit ethanol for a predominant group of frugivorous animals in the tropics, we estimated dietary ethanol exposure for 10 wild primate species using species-specific feeding data and the ripe fruit ethanol concentrations measured in the present study. We focused on peak predicted exposure by examining the 10 days per primate species for which we estimated they would have consumed the highest ethanol concentrations, based on plant species in the diet records. Using mean ethanol concentrations for ripened fruits from each plant species at each sampling location, estimated daily ethanol exposure ranged from near zero (<0.01 g/kg/day) to 0.72 g/kg/day (±1.57 standard deviation (SD)) on average per primate species. The higher value is equivalent to up to 5.1 (±11.0 SD) international standard drinks of 10 g ethanol each per day for a 70 kg human (Fig. 6, table S8). If we instead examine maximum fruit ethanol concentrations, daily exposure estimates were higher for all primate species, ranging from 0.01 (±0 SD) to 15.5 (±10.9 SD) standard drinks per day on average (fig. S5, table S8).

**Fig. 6.**
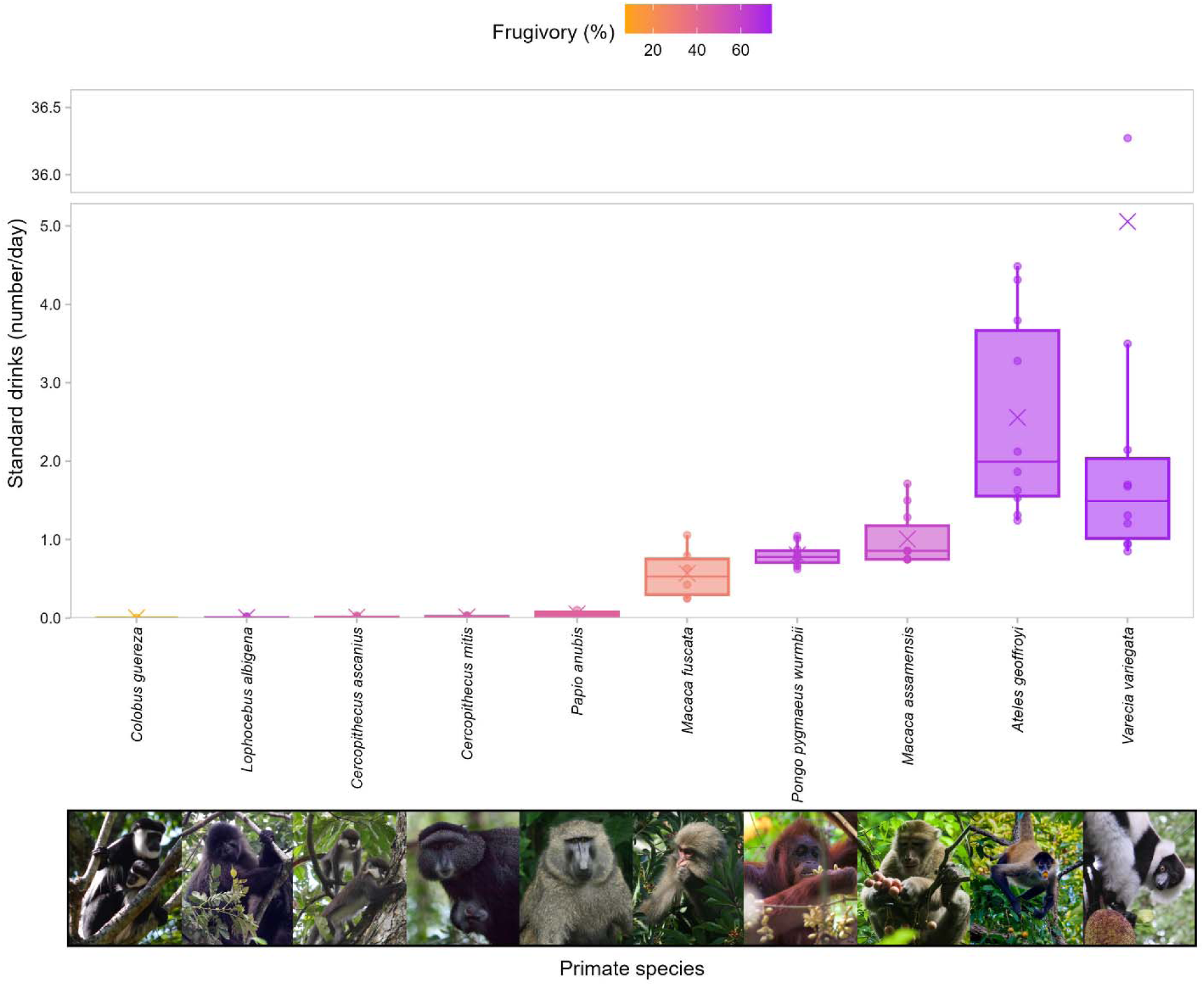
Upper range estimates of daily dietary ethanol exposure in 10 primate species. Ethanol exposure is represented by standard drinks (number/day), equivalent to human consumption of a standard drink adjusted for average adult body mass per primate species. Standard drink estimates are based on primate foraging records and mean ethanol concentration per plant species measured from ripened fruits in this study. Data points represent ethanol consumption per day for an adult primate, with up to the 10 highest values calculated plotted per species. The median and mean standard drinks per species are shown by a horizontal bar and an ‘X’ point, respectively (species are ordered along the abscissa by ascending mean). Average frugivory (% fruit in diet) per species ranges from 7% in *Colobus guereza* (orange) to 74% in *Varecia variegata* (purple) (see table S8 for values and sources). The large outlier for *V. variegata* was verified and thus retained; it relates to a 30-minutes-long foraging event on *Artocarpus heterophyllus* (jackfruit), measured at 0.94 and 1.8% alcohol by volume the day of and after collection, respectively (however, the amount consumed was a mass estimate based on bites taken and could potentially be an overestimate). Photos credits: Photos by Jessica Rothman (*C. guereza*, *Lophocebus albigena*, *Cercopithecus ascanius*, *P. anubis*), Goro Hanya (*M. fuscata*), Rudi Delvaux (*Pongo pygmaeus*), Kittisak Srithorn (*Macaca assamensis*), Nicholas Chapoy (*Ateles geoffroyi*), and Steig Johnson (*V. variegata*), used with permission. *Cercopithecus mitis* photo by Matt Biddulph, licensed under CC BY-SA 2.0 [*cropped from original*], reprinted from (*64*).

Based on the fruit species recorded in their respective diets, and their patterns of feeding behavior, black-and-white ruffed lemurs (*Varecia variegata*), black-handed spider monkeys (*A. geoffroyi*), and Assamese macaques (*Macaca assamensis*) had the highest estimated ethanol exposures. Using mean fruit ethanol concentrations, they were predicted to consume 5.1 (±11.0 SD), 2.6 (±1.3 SD), and 1.0 (±0.4 SD) standard drinks on average, respectively, during the 10 days when ethanol intakes were highest (these values were higher if maximum fruit ethanol values were used; fig. S5, table S8). Bornean orangutans (*Pongo pygmaeus wurmbii*) had an estimated exposure equivalent to just over three-quarters of one standard drink (mean ± SD = 0.8 ± 0.1). Excluding Assamese macaques, other cercopithecoid primates in our dataset had negligible estimated ethanol exposures (Fig. 6).

## DISCUSSION

Through a widespread survey of ethanol concentration in fruits and floral tissue measured across hundreds of angiosperm species, we provide the first large-scale assessment of the ecological distribution of naturally occurring ethanol and the factors shaping its variation. Previous studies have documented ethanol in fruits, but mostly through geographically restricted small-scale surveys (*12*, *24–26*, *35*, *36*), leaving the broader picture of this phenomenon and ecological context unrecognised. Our sampling of 591 angiosperm species across 18 countries and six biogeographical realms shows that ethanol is a widespread component of plant reproductive tissues that occurs across diverse plant taxa and geographic regions. This widespread occurrence supports the hypothesis that ethanol exposure is a common and recurrent feature of the ecology of many frugivorous and nectarivorous animals (*1*, *2*). Our findings indicate that natural ethanol availability emerges from interactions among plant traits, evolutionary history, and environmental conditions.

### Phylogenetic relatedness impacts ethanol concentration

While we found no evidence of a phylogenetic signal in mean or median ethanol concentration, a signal was present in maximum ethanol concentration, indicating that closely related plant species may share characteristics influencing their propensity to accumulate high ethanol concentrations. Families including Anacardiaceae, Arecaceae, Moraceae, and Annonaceae contained multiple species with detectable ethanol and maximum concentrations exceeding 1.5%, whereas other well-sampled families such as Fabaceae and Malvaceae generally showed lower ethanol occurrence and concentrations. Many of the more ethanolic lineages produce fleshy fruits with succulent mesocarps, including berries, drupes, and multiple fruits, whereas comparatively less ethanolic groups often produce dry fruits, capsules, legumes, or fruits with only fleshy arils (*37*, *38*). These differences may influence sugar availability, water content, microbial accessibility, and the suitability of fruit tissues for fermentation. However, distinguishing the relative contributions of plant chemistry, fruit morphology, microbial communities, and dispersal ecology will require targeted comparative studies.

Our finding of phylogenetic structure in maximum ethanol concentration is consistent with previous work from a tropical dry forest in Costa Rica, which identified a strong phylogenetic signal in fruit ethanol concentration among 38 plant species (*35*). The somewhat weaker signal detected in our global dataset may reflect the greater environmental and taxonomic breadth of our sampling, which introduces additional sources of variation that may obscure lineage-level patterns. More broadly, variation in the phylogenetic signal across ecological contexts is common for plant traits, particularly those influenced by interactions between evolutionary history and environmental conditions (*29*, *39*). Our results therefore suggest that ethanol concentration represents a trait with partial evolutionary conservatism, but one that is also strongly shaped by ecological context.

### Environmental conditions shape ethanol availability across tropical ecosystems

Beyond evolutionary history, our results demonstrate that environmental conditions are important predictors of ethanol occurrence and concentration in wild fruits. Ethanol detection was associated with variation in temperature, precipitation, precipitation seasonality, and vegetation density, suggesting that the ecological conditions experienced by plants influence the likelihood that fruits undergo detectable fermentation.

Temperature showed a nonlinear relationship with ethanol occurrence, with detection increasing up to 25°C before declining at higher temperatures. Temperature is expected to have opposing effects on ethanol dynamics. Warmer conditions can enhance microbial activity and accelerate fermentation by increasing yeast growth and metabolic rates (*4*, *27*), potentially increasing ethanol production. However, ethanol is also highly volatile, and elevated temperatures may accelerate evaporation from fruit tissues, or conversion to ethyl esters and acetaldehyde (*40*, *41*); reducing the amount retained at the time of sampling. The decline in ethanol detection at the warmest sites may therefore reflect a balance between enhanced microbial activity and increased ethanol loss, although yeast activity may also decline at very high temperatures.

Vegetation density, measured using NDVI, also predicted ethanol occurrence. More densely vegetated environments may provide conditions favourable for fermentation, including greater humidity, shade, reduced exposure to desiccation, and more stable microbial communities associated with plant surfaces. However, data points at low vegetation densities were relatively sparse, so this result should be treated as preliminary. In addition, vegetation density did not predict mean or maximum ethanol concentration among species, suggesting that these factors may primarily influence whether fermentation occurs rather than the amount of ethanol accumulated once fermentation begins.

Precipitation was the strongest environmental predictor of ethanol concentration. Both mean and maximum ethanol concentrations increased with increasing precipitation, reaching highest values at sites with >3,300 mm annual rainfall. Moisture availability is likely important because yeast activity and microbial persistence depend strongly on water (*6*). The influence of precipitation *seasonality* further suggests that ethanol occurrence may depend not only on average climate but also on temporal patterns of environmental change. Fruits developing in moderately seasonal environments may experience periods that favour both microbial growth and ethanol accumulation, whereas very rainy or extremely dry conditions may constrain fermentation dynamics. Future studies incorporating fine-scale measurements of fruit microclimate, sugar content, microbial communities, and fruiting phenology will be important for identifying the mechanisms linking climate to ethanol variation.

Together, these results provide the first evidence that broad-scale environmental gradients influence natural ethanol availability across angiosperms. Similar to other plant traits shaped by both evolutionary history and climate, such as seed mass, deciduousness, and leaf abscission (*42*, *43*), ethanol concentration appears to emerge from interactions between inherited biological characteristics and local ecological conditions. Changes in vegetation structure and microclimate driven by land use and forest management may therefore alter fruit ethanol availability and dietary exposure among frugivorous animals.

### Fruits collected from the forest floor have higher ethanol concentrations

Our prediction that fruits collected from the forest floor would contain higher ethanol concentrations than fruits collected from the canopy was supported. Accounting for species-level differences, fallen fruits contained greater ethanol concentrations than canopy fruits. This was likely due in part to the larger proportion of fruit in more advanced ripeness stages collected from the ground, but we also found that ripe fruits on the ground had higher ethanol concentrations than ripe fruit collected from the canopy. Several mechanisms may contribute to this result; for example, higher ground humidity, more favorable temperatures, or yeast inoculation and microbial community exposure through fruit fall and contact with soil, leaf litter, insects, and other environmental substrates that may accelerate fermentation processes.

Higher ethanol in ground-sourced fruits has important implications for dietary ethanol exposure among animals. Scrumping fruits from the forest floor is common among terrestrial animals, while some arboreal and volant species also exploit fruit after abscission (*19*, *44–47*). It is also hypothesized that abscission is an adaptation of climacteric fruits (i.e., those that can continue to ripen after separation from the plant), which attract ground-dwelling and olfactorily oriented consumers (*48*). Consequently, dietary ethanol exposure may vary substantially among species depending not only on diet composition but also on foraging strategies and the extent to which animals consume canopy versus ground resources. The ecological importance of fallen fruits may therefore extend beyond energy availability and seed dispersal to include variation in exposure to fermentation products. This result also provides support for hypotheses linking forest-floor foraging with the evolution of increased activity of *ADH4* in the last common ancestor of African apes. Selection acting on this enzyme is posited to have emerged in response to increased consumption of ethanol-containing fallen fruits during terrestrial foraging (*16*, *17*, *49*). Although our study does not directly test evolutionary causation, demonstrating that fallen fruits contain elevated ethanol concentrations provides ecological context consistent with this hypothesis.

### Dietary ethanol exposure across primates as a case study of consumer exposure

Primates are a well-studied taxonomic group with detailed long-term dietary datasets, including for many highly frugivorous species. Estimated daily ethanol intake was generally low for many primates but reached levels equivalent to several human standard drinks in some species when adjusted for body mass. The most frugivorous species in our study, including black-handed spider monkeys and black-and-white ruffed lemurs, had the highest estimated intakes, supporting previous suggestions that dietary specialization may influence ethanol exposure (*13*). Our results are also consistent with previous evidence of ethanol consumption by wild spider monkeys (*24*) and recent estimates of chimpanzee ethanol intake (*25*).

While we did not have sufficient data to incorporate floral resources in dietary estimates, we note that we found ethanol in many flowers and exudates, including in the nectar of *Ravenala madagascariensis*. This plant is a key resource for black-and-white ruffed lemurs (*50*), adding to their high ethanol dietary profile, but also aye-ayes, whose *ADH4* protein suggests a fast ethanol metabolism relative to most other primates, but independently evolved from the *ADH4* modifications underlying faster ethanol metabolism in African great apes and humans (*15*, *16*, *23*, *51*). This supports ongoing research highlighting the ethanolic importance of floral resources (*52*).

Ethanol exposure is unlikely to be restricted to primates. Wild pen-tailed treeshrews (*Ptilocercus lowii*), for example, regularly consume naturally fermented palm nectar, demonstrating recurrent ethanol exposure in a non-primate tropical mammal (*53*). Borneo provides one example of the highly diverse tropical frugivore communities in which numerous sympatric small mammal species, including treeshrews, squirrels and rodents, partition partly overlapping fruit resources, including figs (*54*, *55*). Given the substantial variation in ethanol concentration among plant species, ripeness stages and forest strata documented here, ethanol may represent an overlooked chemical dimension of dietary niche space. Differences among consumers in ethanol detection, avoidance and metabolic tolerance could therefore modify patterns of resource use and dietary overlap within frugivore communities. Because fruits, nectar and other sugar-rich plant resources are consumed by diverse mammals, birds, reptiles and insects, testing this possibility across taxonomically and ecologically diverse consumer communities could reveal whether ethanol contributes to the differentiation of feeding niches and patterns of resource partitioning beyond primates.

### Limitations and future directions

Our sampling was opportunistic and focused on plant species known to occur in the diets of primates and other frugivorous animals. Although this approach allowed us to quantify ethanol availability in ecologically relevant plant resources, it may have biased our dataset toward species accessible to animal consumers and field researchers. Sampling was also uneven among species locations, and seasons, and some plant species, particularly rare taxa and high canopy species, were likely underrepresented. Moreover, because individual fruit species were not sampled continuously across ripening and senescence, we may have missed temporal peaks in ethanol production and thus underestimated maximum ethanol concentrations for some species. Our repeated-sampling thresholds reduced false negatives associated with uneven effort but also excluded many observations from comparative analyses. Additionally, ethanol measurements were constrained by field-based sampling methods, which provide an efficient and scalable method for measuring ethanol across many species and locations, but cannot detect extremely low concentrations, and may underestimate very high concentrations approaching sensor saturation.

Finally, our estimates of consumer exposure were necessarily simplified. The primate dietary analyses relied on existing feeding datasets that varied among sites in sampling intensity and methodology, and were designed to estimate potential upper ranges of exposure rather than daily ethanol consumption. Furthermore, our calculations assume that animals consume fruits in proportion to availability and do not account for behavioural preferences, avoidance, or potential selection for fermented fruits. These variables could usefully be incorporated into future studies. Animals may detect ethanol through olfaction or other sensory cues and actively modify their consumption accordingly (*20–22*).

Regardless of these uncertainties, the widespread occurrence of ethanol in fruits suggests that many consumers must contend with dietary ethanol exposure, highlighting the need to consider ethanol in understanding the ecology and evolution of fruit–consumer interactions. Future research integrating details on plant traits (e.g., sugar, lipid and water content, exocarp thickness, seed number), microbial communities, micro-climate environmental variation, detailed consumer behaviour, and physiological markers of ethanol metabolism will continue to grow our understanding of natural ethanol exposure and evolutionary adaptations across diverse lineages.

Our study provides the first large-scale evaluation of natural ethanol availability in wild plant reproductive tissues. Our results demonstrate that ethanol is a widespread feature of angiosperm fruits and floral resources and the dietary landscape encountered by frugivores. Its occurrence and concentration varied predictably with environmental conditions and forest stratum, while maximum concentrations exhibited phylogenetic structure, indicating that ethanol exposure is neither rare nor randomly distributed across ecological contexts. By linking plant-level ethanol measurements with consumer ecology, we show that natural fermentation represents a pervasive ecological process with potential consequences for frugivore behaviour, physiology, and evolution.

## MATERIALS AND METHODS

### Global sampling of plant reproductive tissues

We sampled angiosperm species included in the diets of primates and other frugivorous animals through a global collaborative network of ecologists and primatologists working at established field sites. Selected field sites encompass high plant diversity and provide opportunities to quantify ethanol availability in plant reproductive tissues relevant to animal diets. Established primate research sites provided a geographically distributed sampling framework because primates are widespread tropical frugivores, and have detailed long-term dietary datasets that enabled subsequent estimates of dietary ethanol exposure.

We sampled wild fruits from 39 locations spanning 18 countries and six major biogeographical realms (Afrotropical, Madagascan, Neotropical, Oriental, Panamanian, and Sino-Japanese (*31*); Fig. 1); the floral tissues of wild angiosperms were also sampled at five of these sampling locations. Sampling locations primarily included natural forests, but we also included cultivated fruits fed on by wild animals (tables S1-S2).

### Quantifying ethanol concentration in fruits and flowers

Researchers opportunistically collected fruits and floral tissues. We defined fruits as both true and accessory fruits (e.g., the cashew apple), including simple, aggregate, and multiple fruit types. Fruits were classified into four ripeness categories (unripe, midripe, ripe, or overripe) by researchers with extensive knowledge of local plant species. Classifications were made using standardized assessments of fruit characteristics including colour, firmness, odour, seed maturity, and evidence of deterioration (*35*). Reliable indicators of ripeness differ among plant taxa, and as such the specific traits used varied among species. Whenever possible, fruits were collected from both canopy and forest-floor locations. Intact and damaged fruits, including those with signs of animal activity (e.g., bite marks, food remnants, or partial consumption; table S1), were sampled and the extent of damage noted. Fruits were considered to be intact if no visible damage resulting in an obvious opening was observed (i.e., tiny punctures, bruises, or scratches). Floral tissue samples were classified as immature buds, mature flowers, or floral exudates (includes nectars or liquid accumulated within floral structures (e.g., bracts)).

Ethanol concentrations were measured following (*35*) using headspace analysis. Samples were weighed, photographed, and placed in sealed plastic bags containing standardized headspace volumes (120 cc of air). Samples were incubated for a minimum of one hour (and ≤12 hours) to allow ethanol to equilibrate between plant tissue and surrounding air. Ethanol standards of known concentration (5 mL each of 0%, 0.5%, 1.0%, and 2.0% ethanol in distilled water) were processed concurrently under equivalent environmental conditions. Headspace air (60 cc) was measured using a Mark V breathalyzer (Alcovisor, Kwun Tong, Hong Kong, China), and resulting blood alcohol content (BAC) measurements were converted to ethanol concentration (% ABV) using sampling-period-specific calibration curves (see supplementary text S1 for further detailed calibration procedures and quality control criteria). Between readings, the breathalyzer was flushed with fresh air through the sensor and all sampling compartments until the machine consistently read 0.000.

Floral tissue samples were not included in statistical analyses because sample sizes were limited (149 samples from 37 angiosperm species); these data are instead presented descriptively. For fruit analyses, we evaluated multiple sampling thresholds requiring repeated sampling of midripe-to-overripe fruits and retained the least restrictive threshold because alternative thresholds produced qualitatively similar results (supplementary text S2).

The repeatedly sampled dataset included 2,575 fruit samples from 211 species across 29 sampling locations, with reliable % ABV ethanol measurements available for 2,454 samples (supplementary text S1 and S2). Species-level ethanol values were calculated as median, mean, and maximum ethanol concentrations, and ethanol occurrence (i.e., presence/absence) was determined based on whether ethanol was detected in at least one reliable measurement for that species across the entire raw dataset. An overview of the steps taken to filter and modify raw data in preparation for each analysis is provided in fig. S1.

### Phylogenetic analysis

To test for phylogenetic signal in fruit ethanol concentration, we generated a phylogenetic tree for the repeatedly sampled dataset (*N* = 211 species; fig. S1) using the ‘GBOTB.extended.WP.tre’ megatree implemented in the phylo.maker2 package (*56*) (Fig. 2). Missing taxa were resolved following procedures described in supplementary text S3.

We estimated Pagel’s λ via maximum likelihood for the repeatedly sampled dataset for three species-level ethanol metrics: mean ethanol concentration, median ethanol concentration, and maximum ethanol concentration (fig. S1). Likelihood ratio tests were used to compare estimated λ values against models where λ was constrained to zero or one (“pgls” function in the “caper” package (*57*); table S9). A Pagel’s λ value of zero is consistent with limited phylogenetic signal, whereas values approaching 1 indicate covariance among species consistent with that expected under Brownian motion evolution (*58*).

We also created a phylogenetic tree and evaluated phylogenetic signal following the above-mentioned methods using the full fruit dataset (i.e., no sampling thresholds and all ripeness stages included; fig. S1), to examine phylogenetic signals across the full range of species (while acknowledging that any significant signal is likely an underestimate of the true value; fig. S4). Our results using the full dataset were similar, but the strength of the detected signal was lower; the lower sampling effort across ripened fruits for the additional 380 plant species in the full dataset likely caused an underestimation of both detected ethanol occurrence and concentrations in many of these species.

### Environmental predictors of fruit ethanol occurrence and concentration

We tested whether environmental conditions predicted ethanol occurrence and concentration using climatic and vegetation variables matched to the geographic coordinates of each sampling location. We extracted annual mean temperature (°C; bio1), annual precipitation (mm; bio12), and precipitation seasonality (coefficient of variation (CV); bio15) from the WorldClim database at a 0.01° resolution (WorldClim version 2.1 (*59*)). We quantified vegetation density using mean NDVI values from MODIS satellite data calculated within a 1 km buffer around sampling locations (MODIS MOD13Q1 version 6 (*60*)) (supplementary text S4).

Using the repeatedly sampled dataset (fig. S1), we aggregated repeated sampling-location-specific measurements from the same plant species to compute estimates of ethanol presence, mean concentration, and maximum concentration per plant species per sampling location. To avoid pseudoreplication, species sampled at multiple locations were represented by a single randomly selected sampling location for environmental analyses (supplementary text S4).

We used zero-inflated generalized linear mixed models (GLMMs; “glmmTMB” function in the “glmmTMB” package (*61*)) to test relationships between ethanol metrics and environmental predictors (table S9). Models included spatial random effects based on sampling-location coordinates. We evaluated linear and second-order polynomial relationships and used corrected Akaike information criterion (AIC_c_)-based model selection to identify the best-supported models.

### Canopy versus forest-floor ethanol concentration

To test whether fruits collected from the tree canopy differed from fruits collected from the ground in ethanol content, we analysed species in our repeatedly sampled dataset for which fruits were repeatedly sampled from both canopy and forest-floor locations and for which ethanol was detected in at least one sample (*N* = 68 species; fig. S1).

We fitted a GLMM (“glmmTMB” function in the “glmmTMB” package (*61*)) to test whether the collection site predicted ethanol concentration while accounting for species-level variation and unbalanced measurement repetition across fruit samples (table S9). Estimated marginal means (EMMs) and a one-sided *Z* test were used to evaluate differences between canopy and forest-floor fruit (supplementary text S5). We repeated these methods using the same GLMM but with the fruit ripeness stage included in the model to test whether there was an interaction between collection site and ripeness stage using a Type III Wald chi-square test. Following the significant finding, we carried out a selection of *post-hoc* two-sided Wald *Z* test contrasts of interest between EMMs of different pairings of collection site and ripeness stage (supplementary text S5).

### Estimating dietary ethanol exposure in primates

To place plant-level ethanol measurements into an ecological context, we estimated the upper range of daily dietary ethanol exposure for 10 wild primate species found at six of our plant sampling locations (Indonesia, Costa Rica, Japan (Yakushima Island), Madagascar, Thailand, Uganda; Fig. 1). For each primate species, we identified feeding days during which individuals consumed fruit from plant species for which we detected ethanol in this study, and estimated daily fruit mass consumed based on feeding duration and plant-species-specific intake rates. Daily ethanol intake was calculated using both mean and maximum measured ethanol concentrations per fruit species per sampling location, based on ripened fruits from the full fruit dataset (fig. S1). Daily ethanol intake was standardized by adult body masses reported in published literature for the primate species and expressed as grams ethanol per kilogram body mass per day (g/kg/day). For each primate species, we selected the 10 foraging days with the greatest predicted ethanol intakes to estimate an upper range of ethanol exposure and converted these to human 10 g international standard drink equivalents following (*25*). Detailed calculations and species-specific inputs are provided in supplementary text S6.

All analyses were conducted in R version 4.6.0 (*62*).

## Acknowledgments

We thank Ansumane Dabo, Evin Murillo Chacon, Saúl Cheves Hernandez, Ronald Lopez Navarro, Mamadu Ba Dabo, Iaia Tawél Camará, Mamadu Cassamá, Razafindravelo Cressant, Mbana Ferdinand, Razafiarisona Jean Clement, Zakamanana Michel Patric, Tsarandro Nomenjanahary Rafidison Benjamin, Rakotoson Emilys Edgarson, Marolahy Jean Piere, Maria Camila Pinzon, Angie Rodriguez Miranda, Kelsey Ellis, Billy Browning, Manuela Bolivar, Leonardo Silva, Peter Irumba, John Rusoke, Hillary Musinguzi, and Miranda Swagemakers for assisting with data collection. We are grateful to the superintendents of Phu Khieo Wildlife Sanctuary for collaboration, Mareike C. Janiak for contributing to early discussions on this topic and Tyler Bonnell for assistance in coding semi-automatic standard curve calculation.

## Permits

This research was conducted under permits from SINAC and the ACG (R-SINAC-ACG-PI-027-18, ACG-PI-037-2019; ACG-064-2023; ACG-PI-033-2023; ACG-PI-011-2024; and ACG-015-2024), the Canada Research Council for Animal Care through the University of Calgary’s Life and Environmental Care Committee (AC19-0167) for research in Costa Rica; the Peruvian Ministry of the Environment (SERFOR; Research Permit #D000116-2021) for research in Peru; the government and the Ministre de la Recherche Scientifique et de l’Innovation Technologique of the CAR and the WWF CAR for research permission and support to collect data in the Dzanga-Sangha Protected Areas in the Central African Republic; the Republic of Indonesia (BRIN Research Permit 130/SIP/IV/FR/8/2022 and permissions from KSDAE, KLHK, BKSDA, BOSF, Indonesia and KPHL Kapuas Kahayan) for research in Indonesia; the Instituto da Biodiversidade e das Areas Protegidas (IBAP) in Guinea-Bissau and Nalu Regulos for research in Cantanhez National Park; Yakushima Forest Ecosystem Conservation Center, Citizens in Inuyama City, and Shizuoka University for research in Japan; the board of ethics of the Secretaría de Medio Ambiente y Recursos Naturales (SEMARNAT, Permit SGPA/DGVS/000041/22) for research in Mexico; the Government of Madagascar, the Kianjavato Commune, and Madagascar Biodiversity Partnership and the Animal Care Committee at the University of Calgary (AC23-0007), and the Institutional Animal Care and Use Committee (IACUC) at Omaha’s Henry Doorly Zoo and Aquarium, and Direction Générale de la Gouvernance Environnementale (Research Permit 441/24/MEDD/SG/DGGE/DAPRNE/SCBE.Re) for permitting research in Madagascar; the National Research Council of Thailand (NRCT) and the Department of National Parks, Wildlife and Plant Conservation of Thailand (DNP-T; Permit 0907.4/15068) for research in Thailand; the Sabah Biodiversity Council (JKM/MBS.1000-2/2 JLD.16 (118), JKM-MBS.1000-2/2, JKM-MBS.1000-2/2(35), JKM-MBS.1000-2/2(63)) for research in Malaysian Borneo; the Ministerio de Ambiente (MAE-DNB-CM-2019-0117, MAATE-DBI-CM-2024-0345) for research in Ecuador; permissions from the Uganda Wildlife Authority and the Uganda National Council for Science and Technology for research in Uganda; permission from the Centre National de la Recherche Scientifique et Tachnologique, Republic of Gabon (AR 030/23/MESRSIT/CENAREST/CG/CST/CSAR) for research in Gabon; permission from the Instituto Federal Goiano, campus Posse (process number 23727.000289.2024-52) for research in Brazil; permissions from the Kingdom of Cambodia Ministry of Environment General Directorate of Natural Protected Areas, Ratanakiri Provincial Department of Environment, and Stung Treng Provincial Department of Environment for research in Cambodia; and the Department of Scientific Services of Gorongosa National Park (PNG/DSCi/C299/2023) for research in Mozambique.

## Funding

Wenner-Gren Foundation (A.D.M., M.A.C., N.J.D.)

Natural Sciences and Engineering Research Council of Canada (RGPIN-2017-03782, A.D.M.; RGPIN/05262-2014, S.E.J.; RGPIN-2025-05384, V.V.V.)

Canada Research Chairs program (950-231257, A.D.M.) Primate Society of Great Britain (A.C.B.)

Fulbright U.S. Student Research Program (W.D.A.)

Transversal Action of Muséum National d’Histoire Naturelle 2020-2021 and 2021-2022 (S.M.)

CNRS long-term SEE-Life funding 2023-2027 (S.M.)

Swiss National Science Foundation (#51NF40_180888 and #310030_185324, L.N.)

L.S.B. Leakey Foundation (E.R.V., W.D.A.)

U.S. National Science Foundation (BCS-0643122 and BCS-1719825, E.R.V.; DDRIG grant no. 2141917, W.D.A.; NSF 1521528, NSF 2217317, and NSF 2223390, J.M.R.)

Rutgers University (E.R.V.)

Rutgers University Center for Human Evolutionary Studies (E.R.V.; Albert Fellows Grant, W.D.A.)

Orangutan Species Survival Plan (W.D.A.)

Global Wildlife Conservation’s Primate Action Fund (W.D.A.) American Philosophical Society Lewis and Clark Fund (W.D.A.) Hunter College (J.M.R.)

DFG (NE 2156/3-1, O.N.)

Calgary Zoological Society (S.E.J.) International Primatological Society (N.H.) Sigma Xi (N.H.)

Asociación para la Conservación de la Cuenca Amazónica (A.C.P., M.W., G.A.E.) Field Projects International (A.C.P., M.W., G.A.E.)

The University of Texas at Austin (A.D.)

Secretaría de Ciencia, Humanidades, Tecnología e Innovación (C.E.R.T.) MEXT Grant-in-Aid (grant no.23H02563, G.H., M.M., F.K., R.K., W.L.) USAid (APS-497-11-000001, E.R.V., S.S.U.A.)

The George Washington University Selective Excellence Fund (E.R.V.) US Fish and Wildlife Service Great Ape Conservation Fund (E.R.V.) The Denver Zoo (E.R.V.)

Programa Institucional de Bolsas de Iniciação Científica do Instituto Federal Goiano (E.B.A.)

Georg-August University of Göttingen (J.O., O.S.)

German Primate Center Leibniz Institute for Primate Research (J.O., O.S.) The Explorers Club Impact Grant (I.R., L.G., M.Anj.)

## Author contributions

Designed the study: A.D.M., K.J.H., M.A.C., J.G.C.

Sampled ethanol in the field: A.A.B.D., A.C.B., A.C.P., A.D., A.D.M., A.N.D., A.U., B.G., C.E.R.T., D.N.R., E.B., E.B.A., E.F., E.R.V., F.K., F.S.A., F.Y.Y.T., G.A.E., G.H., H.R., I.R., J.G.C., J.M.R., J.O., K.J.H., K.W., L.G., L.N., L.T.H.S., L.V.R., M.Anc., M.Anj., M.A.P.L., M.A.B., M.B., M.M., M.W., O.N., O.S., R.K., S.A., S.C., S.E.J., S.M., S.R.M., S.S.U.A., T.R.D., T.G., W.D.A., W.L., X.A.H., Y.K.

Provided primate behavioural data: A.D.M., E.R.V., F.A., G.H., J.M.R., J.O., M.A.P.L., N.H., O.S., S.E.J., W.D.A.

Data curation: J.G.C.

Formal statistical analyses: J.G.C., K.W., A.N.D. Drafted manuscript: J.G.C., A.D.M.

Edited manuscript: J.G.C., A.D.M., A.C.B., K.W., K.J.H., M.A.C., N.J.D., E.R.V., F.A., O.N., S.M., W.D.A.

Funding acquisition: A.D.M., K.J.H., M.A.C.

Project administration, supervision, coordination: A.D.M., K.J.H., M.A.C., J.G.C. All authors reviewed and approved the final manuscript.

## Data, code, and materials availability

All data and code needed to evaluate and reproduce the results in the paper are available at https://github.com/juliacasorso/Ethanol_in_Fruits_and_Flowers. These data and code will be archived in Dryad and Zenodo, respectively, ahead of publication, and the access links will be cited here and DOIs included in the reference list. This study did not generate any new materials.

