## Supplementary Materials for "Pervasive ethanol in the fruits and floral nectars consumed by tropical frugivores"

Julia G. Casorso *et al.*

**This PDF file includes:**

Supplementary Text

Figs. S1 to S5

Legends for tables S1, S2, and S8

Tables S3 to S7 and S9

References

**Other Supplementary Materials for this manuscript include the following:**

Tables S1, S2, and S8

S1. Ethanol measurement calibration and quality control

Ethanol measurements followed the headspace equilibration approach described in detail by (*35*). Samples were incubated in sealed plastic bags alongside ethanol standards of known concentrations. Sampling-period-specific calibration curves were generated from concurrent ethanol standards. Calibration curves were forced through zero and used to convert blood alcohol content (BAC) measurements from the breathalyzer into % alcohol by volume (ABV) values. Samples were measured at least twice in succession whenever possible.

In a small number of cases, reliable standards were not available for a sampling period and we used a standard curve from another period at that sampling location that had similar temperature and humidity values, if available, to calculate % ABV values for samples; otherwise, % ABV values could not be calculated for samples with a BAC value >0, and these sample measurements were used only as evidence of ethanol presence (see points below).

Samples which exceeded the reliable upper detection range of the breathalyzer were remeasured using controlled headspace dilution where possible. Samples with a BAC >0.7 (those nearing ‘breathalyzer saturation’) were re-measured following an air dilution method of 1:2 parts headspace to ambient air, and the resulting % ABV values were multiplied by three to correct for the dilution. Undiluted samples approaching saturation were used only as evidence of ethanol presence if significantly lower than corrected diluted samples (see points below).

To maintain a rigorous dataset, the following sample measures for which the % ABV value could not be reliably obtained were retained solely as evidence of ethanol presence:

- measures with a BAC value >0 for which no reliable standards were available;
- measures from undiluted samples that were significantly lower than corrected measures from diluted samples;
- measures with a BAC value >0 where incubation duration was insufficient for reliable quantification (<1 hour);
- outlying replicate measures for samples with at least three repeat measurements.

Neither % ABV values nor evidence of ethanol presence could be obtained from the following sample measures, which were thus excluded from the full dataset:

- measures with a BAC value equal to zero where incubation duration was insufficient for reliable headspace equilibration (<1 hour);
- outliers that suggested the possibility of human measurement error;
- standard measures that approached ‘breathalyzer saturation’ but that were not diluted and remeasured;
- outlying standard measures that were not within range compared to standard measures from other sampling periods at that sampling location.

If less than three reliable ethanol standards remained following these exclusions, a series of standards from another sampling period with similar temperature and humidity values were substituted, if available, to calculate % ABV values for samples, as described above.

S2. Dataset construction and filtering

Excluding unreliable data points as described in section S1, our raw dataset comprised a total of 8,241 breathalyzer measurements across 4,247 plant samples from at least 591 species (and up to 645 species when including multiple samples of family-level and unknown taxonomic assignments), including multiple measurements from the same species across a total of 39 different sampling locations.

Prior to statistical analyses, which were aggregated at the species level, samples with taxonomic assignments above the level of genus were removed (68 unknown samples, and 11 samples assigned only to taxonomic family). To focus our analyses only on wild plants, we removed any provisioned food items or fruits obtained from local markets. Floral tissue samples were not included in statistical analyses because sample sizes were limited (149 samples from 37 angiosperm species across five sampling locations). The resulting full fruit dataset comprised 8,120 breathalyzer measurements across 4,173 fruit samples, 591 species, and 39 sampling locations.

Species-level comparative analyses required repeated sampling because species with greater sampling effort had higher probabilities of ethanol detection. To reduce the likelihood of false negatives, we evaluated three minimum sampling thresholds: (1) at least three midripe, ripe, or overripe fruits sampled per plant species; (2) at least three ripe or overripe fruits per species; and (3) at least three ripe fruits per species. All thresholds produced qualitatively similar results, and the least restrictive threshold was retained. Unripe fruits were excluded from analyses because ethanol was rarely detected at this stage (93% of unripe samples had 0% ABV). The final, repeatedly sampled dataset used for analyses consisted of 5,197 breathalyzer measurements from 2,575 fruit samples, 211 species, and 29 sampling locations. Reliable % ABV measures were obtained from 2,454 of these fruit samples, with the remainder of samples retained solely as evidence of ethanol presence. Data filtering steps are visualized in fig. S1.

S3. Phylogenetic tree construction

We used the GBOTB.extended.WP.tre mega-tree (Jin & Qian, 2022), which comprises 72,750 species of vascular plants across over 480 families and follows the World Plant database for nomenclature (*65*), to construct a phylogenetic tree of our fruit species from the repeatedly sampled dataset (“phylo.maker*”* function in the “V.PhyloMaker2” package (*56*)). For species or genera absent from the mega-tree, accepted synonyms or basionyms were substituted where available. Otherwise, a missing species or genus was attached as a sister taxon to a closely related species or genus in the mega-tree (using the ‘species.relative’ or ‘genus.relative’ columns in our species list, as described in (*66*)). If there was more than one option of an equally closely related taxon, the first alphabetical option present in the mega-tree was chosen, while ensuring the distance of relation between the chosen relative and the missing taxon was equal to any other equally-related-taxa already present in our species list (determined using the Open Tree of Life (*67*)).

Samples identified only to the level of genus were attached as sister taxa to their respective genera. Such samples were included only when it could be determined they were distinct from other known species within the same genus that were present in our species list.

We separately followed these steps to resolve missing taxa and coarse taxonomic resolution on the list of species from the full fruit dataset for which we had reliable % ABV values (*N* = 583 species) to create a phylogenetic tree for use in analyzing our full range of species (while acknowledging that any significant results are likely underestimates).

S4. Environmental variable extraction and modelling

For quantification of environmental predictors, we prioritized open-source datasets over localized, snapshot field temperature measurements to enhance transparency and facilitate reproducibility in future research. We extracted climatic information from the WorldClim database version 2.1, which contains high-resolution global climate data from 1970 to 2000 (*59*), and extracted vegetation density from the 2023 satellite data layer of MODIS MOD13Q1 version 6 (*60*).

We selected several predictor variables to meaningfully capture environmental contrasts relevant to plant growth and fruiting dynamics across the diverse ecosystems and biogeographical regions included in our study. To represent annual climatic trends, we selected (1) annual mean temperature (°C; bio1), or the yearly average of all average monthly temperatures, and (2) annual precipitation (mm; bio12), or the sum of total monthly precipitation values across a year (*68*). To represent seasonality, we selected precipitation seasonality (coefficient of variation (CV); bio15), or the variation in monthly precipitation, calculated as the ratio of the standard deviation (SD) of monthly total precipitation to the mean monthly total precipitation and expressed as a percentage (larger values represent greater variability). CV values that exceed 100%, represent regions that have very little rainfall such that any variation causes an extreme percentage (e.g., deserts) or climatic anomalies (typically confined to coastal areas and islands (*68*)). Lastly, we quantified overall vegetation density at sampling locations based on the normalized difference vegetation index (NDVI), computed as mean and one SD of all repeated measures in a 1 km buffer around sampling locations. We further characterized all sampling locations by biogeographical realm (Afrotropical, Madagascan, Neotropical, Oriental, Panamanian, Sino-Japanese) using the zonation proposed by (*31*) (though this variable was not included in any of the resulting best-supported models; table S9).

The repeatedly sampled dataset was aggregated at both the level of plant species and sampling location to create unique site- and species-specific ethanol metrics. ‘Duplicates’ were removed if the same species was sampled at multiple sites, using a ‘random’ procedure whereby samples from all but one site were removed via a duplicate omission function during data processing (resulting in the exclusion of three sites, and no reduction in the number of species). We evaluated the influence of all environmental predictors on the following metrics: (1) ethanol presence, or the percentage of species at a sampling location that had ethanol detected in at least one reliable fruit measured across the entire raw dataset, and (2) ethanol concentration, calculated as either (a) mean or (b) maximum ethanol concentration per species per sampling location. Models including ethanol concentration were run only on non-zero mean and maximum values (resulting in the exclusion of an additional four sites and 78 species). For each metric, we fitted several generalized linear mixed models (GLMMs) with combinations of linear predictors and second order orthogonal polynomial terms (assuming that unimodal relationships are likely because both relatively low and high values of environmental variables may be less favorable to ethanol production; table S9). We then used variable selection based on the Akaike information criterion corrected for sample size (AIC_c_) to identify the most parsimonious models for prediction.

S5. Canopy versus Ground modelling and *post-hoc* contrasts

To compare ethanol between canopy- and ground-collected fruits, we focused on a subset of the repeatedly sampled dataset comprised of species that had (1) fruits sampled from both the canopy and forest floor (minimum of one fruit per category) and (2) had ethanol detected in at least one reliable sample across the entire raw dataset (*N* = 68 species). We evaluated the influence of collection site (ground, canopy) on ethanol proportion by fitting a GLMM with the following random effects: (1) fruit species name; (2) fruit intactness when measured (categorized as intact, non-intact, or unknown); (3) sample ID; and (4) fruit ID. The inclusion of sample ID accounted for uneven breathalyzer measurement replication across samples, while the inclusion of fruit ID accounted for uneven numbers of samples between individual fruits (e.g., a fruit cut in half with both halves measured as separate samples, compared to a different fruit only sampled in its original state). We compared the model both with and without zero-inflation (using the Smithson and Verkuilen transformation on proportion ethanol to remove zeroes and enable a beta-logit distribution with the non-zero inflated model (*69*)), finding that the latter was most supported according to the AIC_c_. We executed a one-sided *Z* test on the estimated marginal means (EMMs) from the non-zero-inflated model, based on our alternative hypothesis that ethanol in ground-collected fruits would be greater than that in canopy-collected fruits.

We repeated this analysis using the same model but with fruit ripeness stage (midripe, ripe, overripe) included as an interaction term and executing a Type III Wald chi-square test on the model. Upon finding a significant interaction, we extracted EMMs for all possible pairings of collection site and ripeness stage, and executed two-sided Wald *Z* test contrasts on the following pairs: (1) Ground-Midripe versus Plant-Midripe; (2) Ground-Ripe versus Plant-Ripe; (3) Ground-Overripe versus Plant-Overripe; (4) Ground-Midripe versus Ground-Ripe; (5) Ground Ripe versus Ground-Overripe; (6) Plant-Midripe versus Plant-Ripe; and (7) Plant-Ripe versus Plant-Overripe. We used the Holm-Bonferroni correction to adjust *P* values for all seven contrasts. After finding that the sixth contrast was non-significant and the seventh contrast was significant, we added an eighth contrast comparing the average of Plant Midripe & Plant Ripe versus Plant Overripe. We evaluated all eight contrasts together, again adjusting *P* values with the Holm-Bonferroni correction now for all eight contrasts (table S7).

S6. Primate dietary exposure calculations

Dietary exposure estimates were based on feeding records from 10 wild primate species across six long-term field sites used as plant sampling locations (table S8). Because field datasets differed in sampling duration and resolution, analyses focused on upper ranges of dietary ethanol exposure, whereby we selected and compared a subset of days for which we calculated the highest daily intake rates per primate species.

To estimate ethanol intake, we first organized foraging records by individual focal ID (focusing only on adults), date, and plant species consumed. The foraging duration (number minutes spent feeding) for each record was multiplied by the primate-species-specific intake rate for that plant species (grams wet weight per minute) to estimate mass of fruit consumed (grams wet weight). We then multiplied the estimated fruit mass consumed by a) the mean and b) the maximum ethanol concentration we recorded for that fruit species at that sampling location to estimate the amount of ethanol consumed in grams. To maximize the number of plant species and foraging records included in the primate diet analyses, we used all midripe, ripe, and overripe fruits in the full fruit dataset (i.e., no minimum sampling thresholds applied and excluding unripe fruits) to calculate sampling-location-specific mean and maximum ethanol concentrations per plant species. Next, estimates of ethanol consumption were summed across all records per day per individual focal ID to get an estimate of individual daily mass ethanol intake (grams per day). We divided each intake value by the average adult weight for that primate species to get a body mass-specific dosage (grams per kilogram per day; see table S8 for body mass values and sources). Finally, to compare these dosages to that of a single standard drink ingested by a human, we divided them by a “human conversion factor” (i.e., 10 g ethanol in a 70 kg human = 0.14 g/kg (*25*)) to estimate the number of standard drinks consumed per day.

For each primate species, we selected the ten highest daily intake values (separately for mean- and maximum-ethanol-based values) and calculated the mean (± SD) and median (presented in Fig. 6, fig. S5, and table S8). For three species (*Colobus guereza*, *Papio anubis*, and *Macaca fuscata*), only one, two, and six daily standard drink values, respectively, could be calculated due to availability of overlapping foraging data and ethanol measurements for plant species.

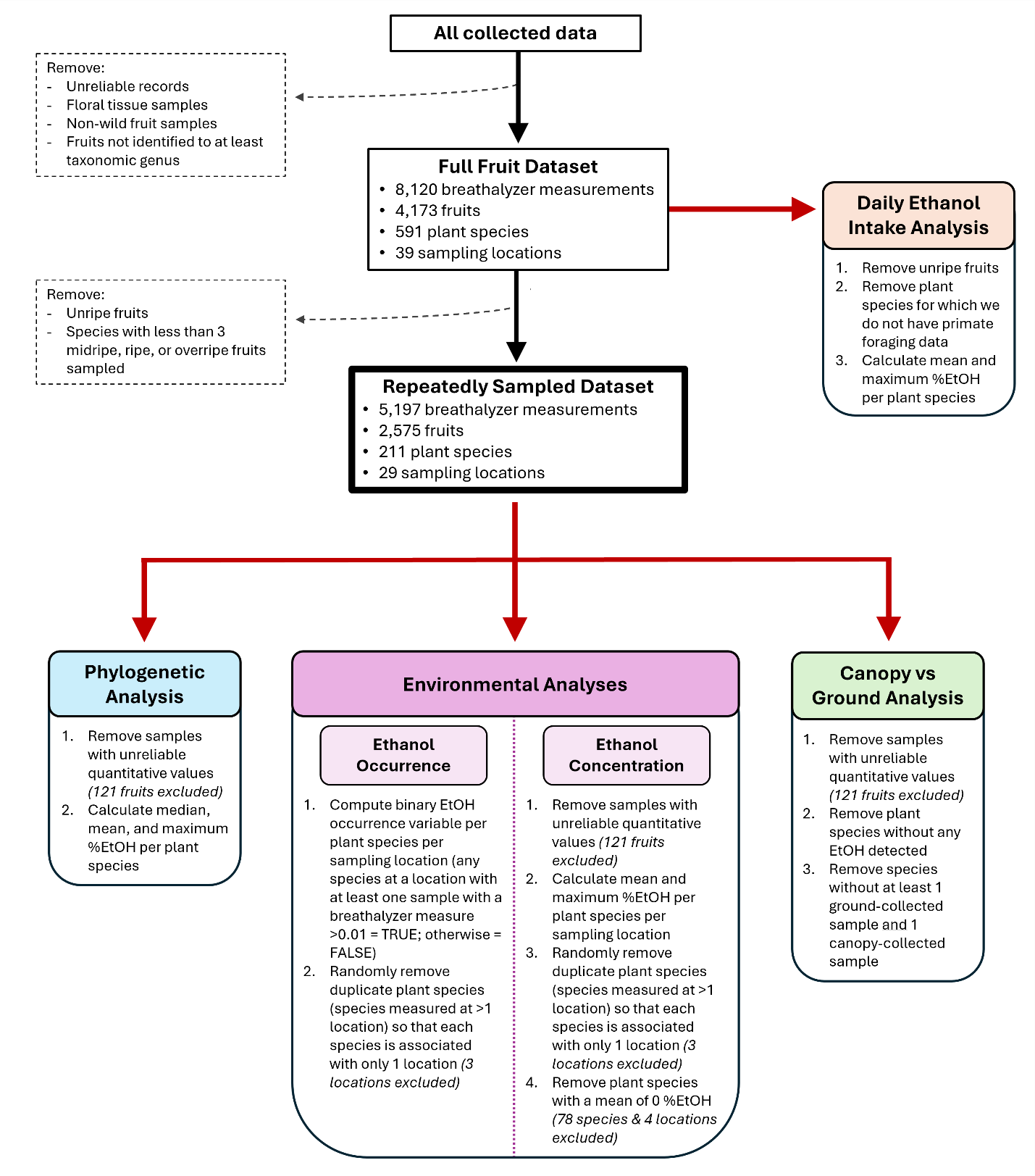

Fig. S1. Flowchart visualizing steps taken to filter and modify the raw dataset prior to executing different analyses in this study. The daily ethanol intake analysis takes ethanol concentrations from the full fruit dataset, rather than the repeatedly sampled dataset, in order to maximize the number of plant species included and therefore maximize usage of available primate foraging data. %EtOH refers to ethanol concentration with units % alcohol by volume (ABV). See supplementary text for further details.

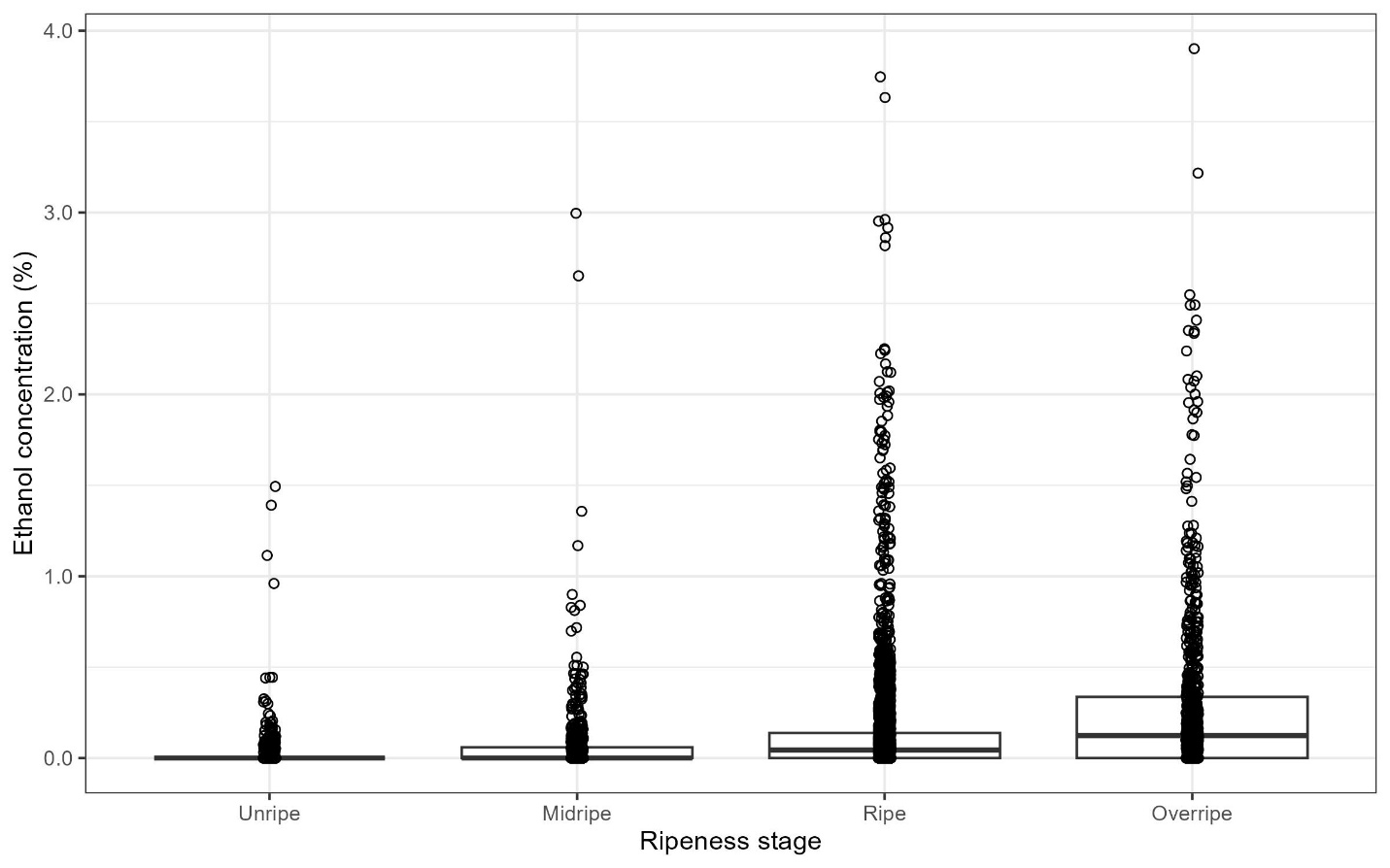

Fig. S2. Distribution of ethanol concentrations in fruits of different ripeness stages. Data are from all 272 fruit species with reliable ethanol concentration (% alcohol by volume (ABV)) values that had ethanol detected in at least one reliable measure in our dataset (comprising a total of 5,683 breathalyzer measurements), separated by ripeness stage (unripe: *N* = 718 samples; midripe: *N* = 404 samples; ripe: *N* = 1586 samples; overripe: *N* = 718 samples). Each data point represents a single breathalyzer measurement, converted to % ABV, with an introduced horizontal error of magnitude 0.02% ABV to improve visualization (darker areas indicate greater overlap of points). Ripeness stages are ordered along the abscissa by increasing median value, which is denoted by a horizontal bar in each box.

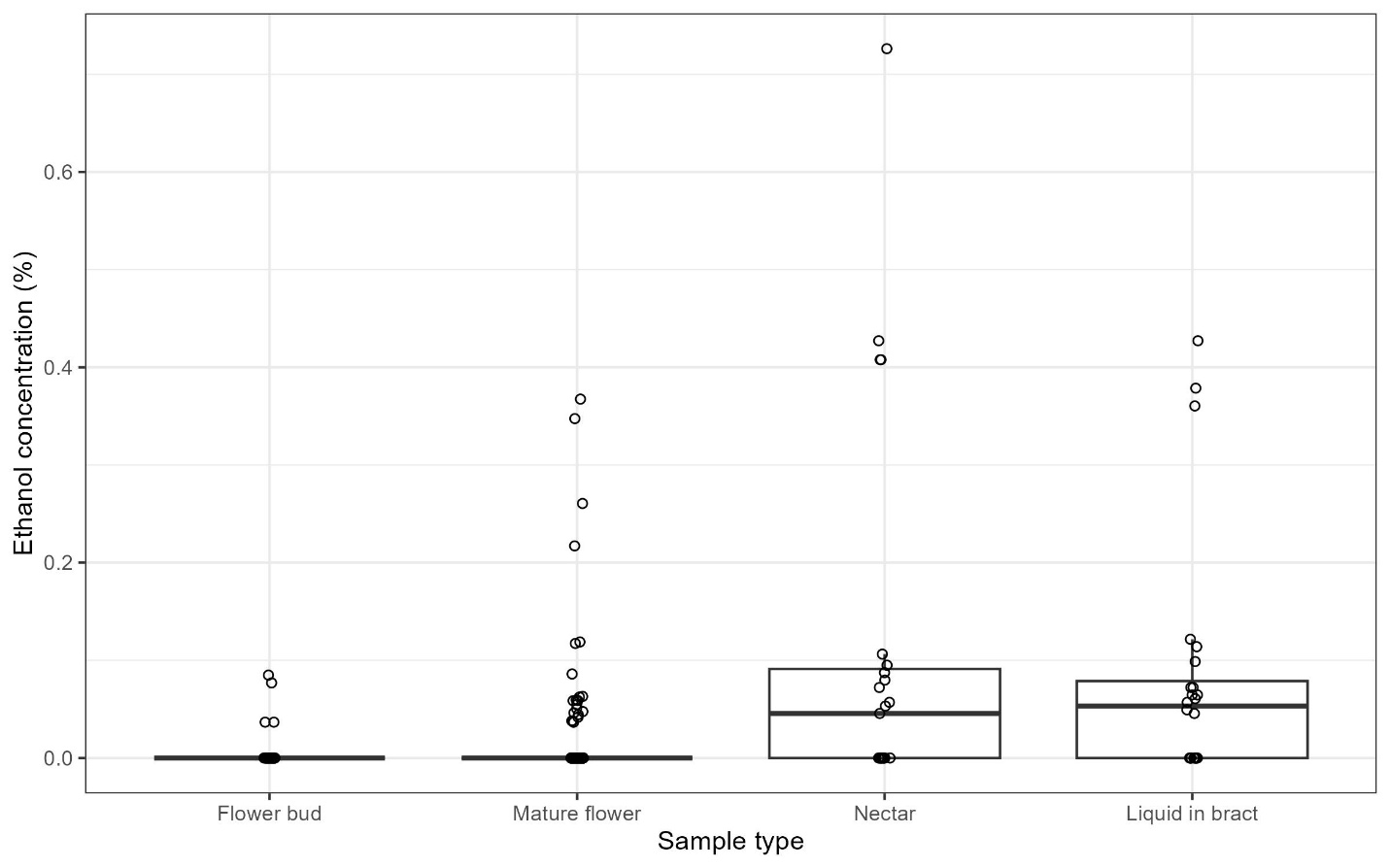

Fig. S3. Distribution of ethanol concentrations across floral tissue samples. Data are from all 37 species with floral tissue samples (comprising a total of 274 breathalyzer measurements), separated by sample type (flower bud: *N* = 20 samples; mature flower: *N* = 101 samples; nectar: *N* = 12 samples; liquid in bract: *N* = 14 samples). Each data point represents a single breathalyzer measurement, converted to ethanol concentration (% alcohol by volume), with an introduced horizontal error of magnitude 0.02% to improve visualization (darker areas indicate greater overlap of points). Sample types are ordered along the abscissa by increasing median value, which is denoted by a horizontal bar in each box. Nectar and liquid accumulated inside bracts were sampled only from the traveller’s tree, *Ravenala madagascariensis*, which has been hypothesized to be a possible fermented nectar source for frugivores (*23*). Flower samples that contained a mixture of sample types (i.e., combination of flower buds and mature flowers (*N* = 7 samples), combination of mature flowers and unripe fruits (*N* = 1 sample)) were assigned to the mature flower category for the plot.

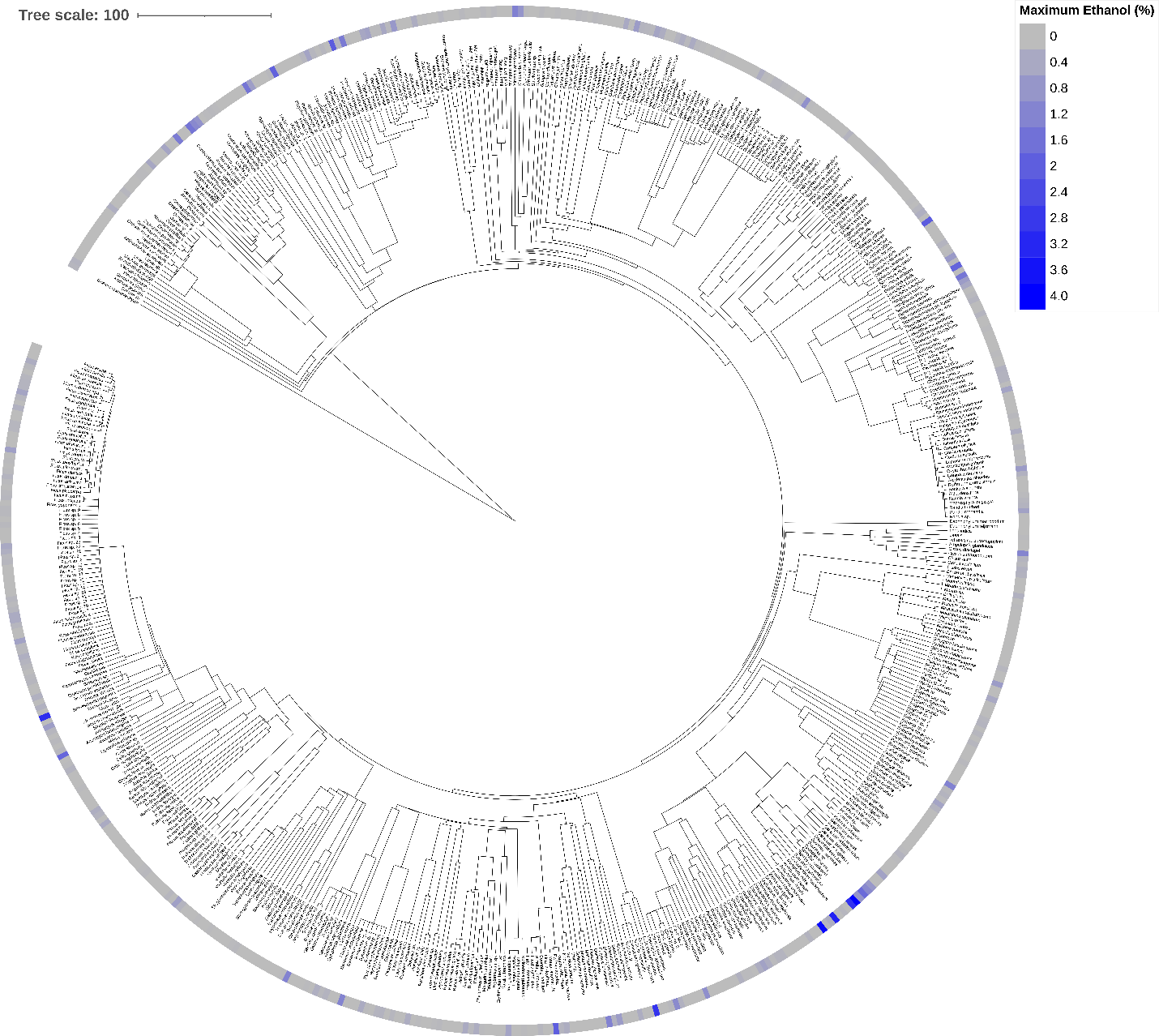

Fig. S4. Phylogenetic tree of all fruit species sampled in this study and their highest ethanol concentration detected. All 583 fruit species (classified to at least taxonomic genus) sampled across all ripeness stages are included in this tree. The maximum ethanol concentration (% alcohol by volume) detected per species is visualized along a colour gradient, with darker colours indicating higher values. Tree scale reflects a distance of 100 Ma. This tree was generated using the mega-tree ‘GBOTB.extended.WP.tre’ in V.PhyloMaker2 (*56*) in R version 4.6.0 (*62*), and visualized with Interactive Tree of Life (iTOL) version 6 (*63*).

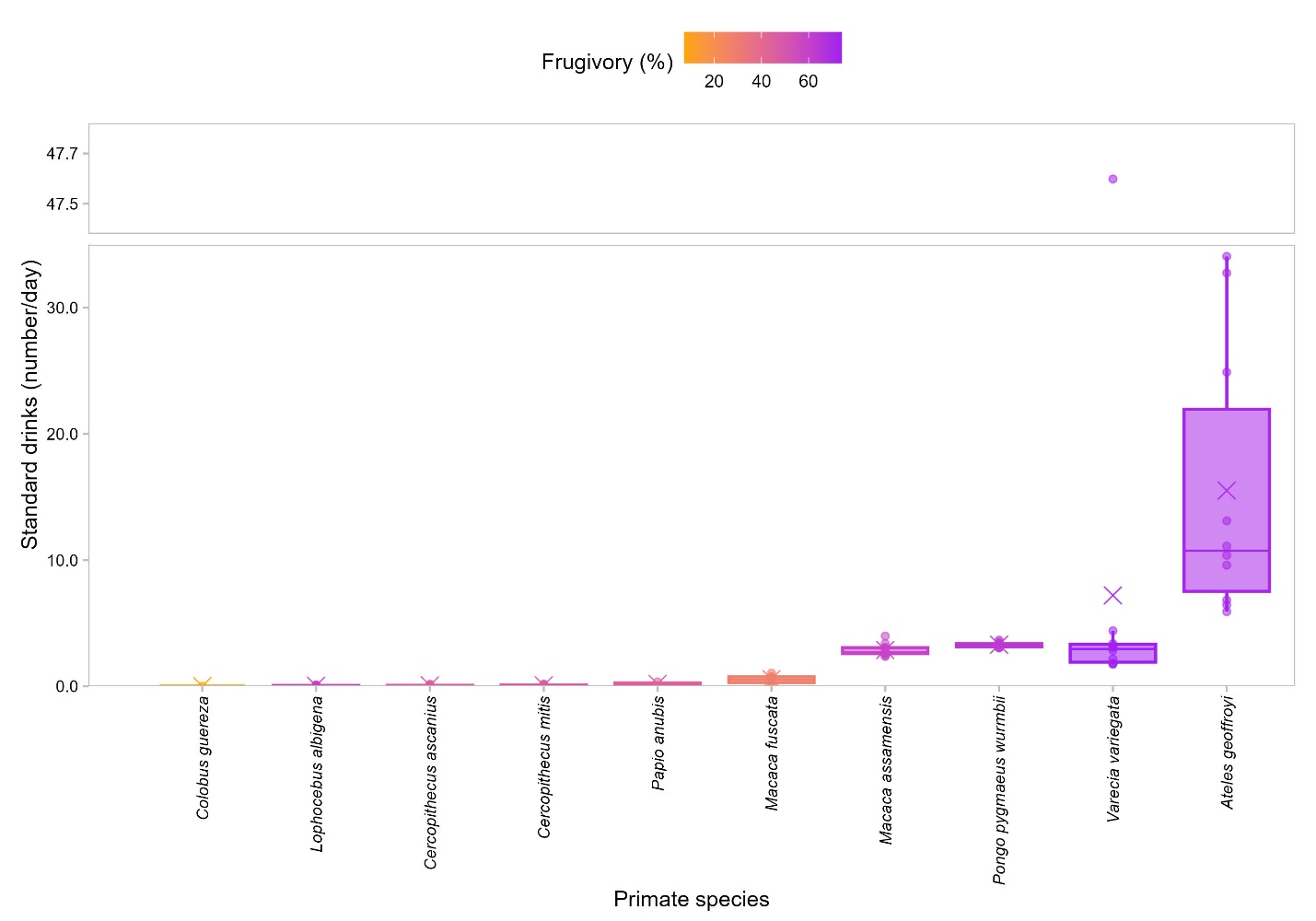

Fig. S5. Upper range estimates of daily dietary ethanol exposure in 10 primate species, based on maximum plant ethanol concentrations. Ethanol exposure is represented by standard drinks (number/day), equivalent to human consumption of a standard drink adjusted for average adult body mass per primate species. Standard drink estimates are based on primate foraging records and maximum ethanol concentration per plant species measured from ripened fruits in this study. Data points represent ethanol consumption per day for an adult primate, with up to the 10 highest values calculated plotted per species. The median and mean standard drinks per species are shown by a horizontal bar and an ‘X’ point, respectively (species are ordered along the abscissa by ascending mean). Average frugivory (% fruit in diet) per species ranges from 7% in *Colobus* *guereza* (orange) to 74% in *Varecia* *variegata* (purple) (see table S8 for values and sources). Note *C. guereza*, *Papio* *anubis*, and *Macaca* *fuscata* distributions include only one, two, and six standard drink values, respectively, due to availability of data. The large outlier for *V. variegata* was verified and thus retained; it relates to a 30-minute-long foraging event on *Artocarpus* *heterophyllus* (jackfruit), measured at 0.94 and 1.8% ABV the day of and after collection, respectively (however, the amount consumed was a mass estimate based on bites taken and could potentially be an overestimate).

Table S1. Ethanol concentrations and sample sizes of all 591 species included in the full fruit dataset. Each fruit species is listed along with the following: taxonomic order and family; mean and maximum ethanol concentrations (EtOH_mean_ and EtOH_max_, respectively; % alcohol by volume (ABV)) across all fruits of all ripeness stages; total number of breathalyzer measurements taken across all ripeness stages and samples (*N*_measures_); total number of fruits that were sampled (*N*_fruits_); number of individual midripe, ripe, and overripe fruit samples with reliable % ABV ethanol measurements (*N*_ripenedfruits_). We also list samples with evidence of animal activity (number of fruits in parentheses), whereby researchers documented an invertebrate, bird, and/or mammal interacting with the sample prior to collection (e.g., sniffing, handling, biting, etc.); a slash between two groups denotes either-or. Note fruits with animal activity were not necessarily non-intact. Lastly, we list the biogeographical realm(s) from which samples of each species were collected in this study (following zonation in (*31*)). Note many species can occur in multiple biogeographical realms, but were not sampled across all. Some species with detectable levels of ethanol have an EtOH_mean_ and EtOH_max_ of 0.00%, as any non-zero breathalyzer measurements were obtained in instances where a % ABV value could not be reliably obtained.

________________________

__________________________

*Measurable ethanol was detected in at least one reliable fruit sample for this species across the entire raw dataset.

†This species was included in the repeatedly sampled dataset (i.e., at least three samples per species, excluding unripe fruits) used for analyses.

Table S2. Ethanol concentrations and sample sizes of all 37 species of flowers and floral exudates sampled in our study. Each plant species is listed along with the following: taxonomic order and family; mean and maximum ethanol concentrations (EtOH_mean_ and EtOH_max_, respectively; % alcohol by volume (ABV)); whether ethanol was detected in fruits of the same species, if applicable (Fruit EtOH_presence_; TRUE = detected ethanol in at least one reliable fruit measurement across our entire dataset); number of individual flower/floral exudate samples (*N*_samples_); and the biogeographical realm from which samples of each species were collected in this study (following zonation in (*31*)). Note many plant species can occur in multiple biogeographical realms, but were not sampled across all. Note *Prismatomeris tetrandra* had detectable levels of ethanol but its EtOH_mean_ and EtOH_max_ concentrations are 0.00%; this is because ethanol was detected only in measurements where a % ABV ethanol value could not be reliably obtained (i.e., retained for evidence of ethanol presence/absence).

__________________________

*Measurable ethanol was detected in at least one reliable floral tissue sample for this species across the entire raw dataset.

†At least three floral tissue samples were measured for this species.

‡Fruits from this species were included in the repeatedly sampled dataset used for analyses.

Table S3. Complete results of the analysis of the impact of environmental predictors on fruit ethanol occurrence. Results are from the binomial (logit link) generalized linear model of first and second order polynomial relationships between species-level ethanol occurrence (presence/absence) and mean annual temperature (‘Temperature’), annual precipitation (‘Precipitation’), precipitation seasonality (‘Seasonality’), and mean Normalized Difference Vegetation Index (‘NDVI’) for the repeatedly sampled dataset (i.e., at least three samples per species, excluding unripe fruits, *N* = 211 species). This model was selected based on the corrected Akaike information criterion (AIC_c_) as the most parsimonious model for prediction. Table columns represent the fixed effect variables, estimated coefficient, standard error (SE) of the coefficient, *Z* value (estimate / SE), and *P* value associated with the *Z* value.

| Fixed effect | Estimate | SE | *Z* | *P* |
| --- | --- | --- | --- | --- |
| (Intercept) | 0.733 | 0.177 | 4.132 | 3.590e-05 * |
| Temperature (linear) | 11.425 | 4.003 | 2.854 | 0.004 * |
| Temperature (quadratic) | -9.107 | 3.970 | -2.294 | 0.022 * |
| Precipitation (linear) | 12.198 | 3.906 | 3.123 | 0.018 * |
| Precipitation (quadratic) | -1.480 | 3.451 | -0.429 | 0.668 |
| Seasonality (linear) | 7.592 | 3.167 | 2.398 | 0.017 * |
| Seasonality (quadratic) | -2.636 | 3.631 | -0.726 | 0.468 |
| NDVI (linear) | 3.906 | 2.552 | 1.531 | 0.126 |
| NDVI (quadratic) | -5.178 | 2.578 | -2.008 | 0.045 * |

__________________________

*Statistical significance (α = 0.05).

Table S4. Complete results of the analysis of the impact of environmental predictors on fruit mean ethanol concentration. Results are from the Gamma (log link) generalized linear model for first and second order polynomial relationships between species-level mean ethanol concentration (proportion alcohol by volume) and mean annual temperature (‘Temperature’), annual precipitation (‘Precipitation’), and precipitation seasonality (‘Seasonality’) for the repeatedly sampled dataset (i.e., at least three samples per species, excluding unripe fruits, *N* = 211 species). This model was selected based on the corrected Akaike information criterion (AIC_c_) as the most parsimonious model for prediction, across both mean and maximum ethanol concentration. Table columns represent the fixed effect variables, estimated coefficient, standard error (SE) of the coefficient, *Z* value (estimate / SE), and *P* value associated with the *Z* value.

| Fixed effect | Estimate | SE | *Z* | *P* |
| --- | --- | --- | --- | --- |
| (Intercept) | -6.733 | 0.097 | -69.159 | < 2.000e-16 * |
| Temperature (linear) | -2.093 | 1.749 | -1.197 | 0.231 |
| Temperature (quadratic) | -0.744 | 1.283 | -0.580 | 0.562 |
| Precipitation (linear) | 5.395 | 1.485 | 3.633 | 2.810e-4 * |
| Precipitation (quadratic) | 0.216 | 1.486 | 0.146 | 0.884 |
| Seasonality (linear) | 3.357 | 1.283 | 2.616 | 0.009 * |
| Seasonality (quadratic) | -3.396 | 1.889 | -1.798 | 0.072 |

__________________________

*Statistical significance (α = 0.05).

Table S5. Complete results of the analysis of the impact of environmental predictors on fruit maximum ethanol concentration. Results are from the Gamma (log link) generalized linear model for first and second order polynomial relationships between species-level maximum ethanol concentration (proportion alcohol by volume) and mean annual temperature (‘Temperature’), annual precipitation (‘Precipitation’), and precipitation seasonality (‘Seasonality’) for the repeatedly sampled dataset (i.e., at least three samples per species, excluding unripe fruits, *N* = 211 species). This model was selected based on the corrected Akaike information criterion (AIC_c_) as the most parsimonious model for prediction, across both mean and maximum ethanol concentration. Table columns represent the fixed effect variables, estimated coefficient, standard error (SE) of the coefficient, *Z* value (estimate / SE), and *P* value associated with the *Z* value.

| Fixed effect | Estimate | SE | *Z* | *P* |
| --- | --- | --- | --- | --- |
| (Intercept) | -5.462 | 0.092 | -59.375 | < 2.000e-16 * |
| Temperature (linear) | 1.491 | 1.669 | 0.893 | 0.372 |
| Temperature (quadratic) | 0.184 | 1.342 | 0.137 | 0.891 |
| Precipitation (linear) | 4.004 | 1.410 | 2.839 | 0.005 * |
| Precipitation (quadratic) | 0.999 | 1.430 | 0.698 | 0.485 |
| Seasonality (linear) | 2.992 | 1.197 | 2.500 | 0.012 * |
| Seasonality (quadratic) | -2.909 | 1.827 | -1.592 | 0.111 |

__________________________

*Statistical significance (α = 0.05).

Table S6. Complete results of the analysis of the impact of collection site and ripeness stage on fruit ethanol concentration. Results are from a Type III Wald chi-square test on the Beta (logit link) generalized linear mixed model that tested for an interaction between collection site (ground, canopy) and ripeness stage (midripe, ripe, overripe). Data come from the repeatedly sampled dataset (i.e., at least three samples per species, excluding unripe fruits), further filtered for species that had (a) ethanol detected in at least one reliable measure in our entire dataset, and (b) at least one fruit sampled from both the ground and the canopy (*N* = 68 species). Table columns represent the fixed effect variables in the model, estimated Wald chi-square test statistic (χ^2^), degrees of freedom (df), and *P* value associated with the chi-square value. The term ‘x1:x2’ denotes an interaction effect between the two independent variables (‘x1’ and ‘x2’) in the model.

| Fixed effect | χ^2^ | df | *P* |
| --- | --- | --- | --- |
| (Intercept) | 819.453 | 1 | < 2.200e-16 * |
| CollectionSite | 1.687 | 1 | 0.194 |
| RipenessStage | 68.663 | 2 | < 1.231e-15 * |
| CollectionSite:RipenessStage | 14.084 | 2 | 8.743e-04 * |

__________________________

*Statistical significance (α = 0.05).

Table S7. Results of *post hoc* contrasts investigating the interaction between collection site and ripeness stage from the canopy versus ground analysis. Each row represents an estimated marginal mean two-sided Wald *Z* test performed on the log odds ratio scale for a selected combination of collection site (ground, canopy) and ripeness stage (midripe, ripe, overripe) levels. Table columns represent: the contrast test number; a description of the group levels compared per contrast; estimated odds ratio (on the response scale); standard error (SE) of the coefficient; *Z* value (log odds scale estimate / SE); *P* value associated with the *Z* value (all *P* values are adjusted for eight multiple contrasts via the Holm-Bonferroni correction); and 95% confidence intervals (CI) back-transformed from the log odds ratio scale, listed as ‘[lower limit, upper limit]’. All contrasts had infinite degrees of freedom.

| Contrast number | Description | Odds ratio | SE | *Z* | Corrected *P* | 95% CI |
| --- | --- | --- | --- | --- | --- | --- |
| C1 | Ground versus canopy (midripe only) | 1.188 | 0.157 | 1.299 | 0.376 | [0.827, 1.705] |
| C2 | Ground versus canopy (ripe only) | 1.438 | 0.100 | 5.198 | < 1.000e-04 * | [1.188, 1.740] |
| C3 | Ground versus canopy (overripe only) | 0.789 | 0.122 | -1.533 | 0.376 | [0.518, 1.204] |
| C4 | Midripe versus ripe (ground only) | 0.701 | 0.070 | -3.546 | 1.600e-03 * | [0.534, 0.922] |
| C5 | Ripe versus overripe (ground only) | 0.545 | 0.049 | -6.701 | < 1.000e-04 * | [0.426, 0.698] |
| C6 | Midripe versus ripe (canopy only) | 0.849 | 0.093 | -1.499 | 0.376 | [0.630, 1.144] |
| C7 | Ripe versus overripe (canopy only) | 0.299 | 0.042 | -8.549 | < 1.000e-04 * | [0.203, 0.440] |
| C8 | Average of midripe & ripe, versus overripe (canopy only) | 0.276 | 0.039 | -9.064 | < 1.000e-04 * | [0.187, 0.407] |

__________________________

*Statistical significance (α = 0.05).

Table S8. Upper range estimates of daily dietary ethanol exposure in ten primate species, based on either mean or maximum ethanol concentration in fruits. Table columns list each primate species’ taxonomic and common name, the study site and country from which data was collected (equivalent between the foraging and ethanol datasets), their average adult body weight (kg), their average frugivory (% of diet made up of fruit), and measures of daily ethanol consumption based on either mean or maximum ethanol concentration per fruit species measured from ripened fruits. Ethanol consumption was calculated per foraging event on a fruit species by multiplying the estimated mean wet weight (g) consumed by either the mean or maximum ethanol concentration for that fruit species. Dosages were summed together across all fruit species eaten in a day to obtain an estimated daily ethanol consumed (g/day). These values were divided by the average adult body weight (kg) per primate species to obtain a body mass-specific dosage (g/kg/day). Lastly, these dosages were divided by a “human conversion factor” of 0.14 g/kg (i.e., 10 g ethanol in 1 standard drink consumed by a 70 kg human) to obtain values equivalent to the consumption of standard drinks (number/day). All consumption values in the table are listed as mean ± standard deviation (SD), calculated from the ten highest individual daily consumption values computed (however, for *Colobus guereza*, *Papio anubis*, and *Macaca fuscata*, values were calculated from only one, two, and six daily consumption values, respectively, due to availability of data).

__________________________

*Average adult body weight based on (*70*) for *Ateles geoffroyi*; (*71*) for *Cercopithecus ascanius*, *Cercopithecus mitis*, *C. guereza*, *Lophocebus albigena* (reported as *Cercocebus albigena*), and *P. anubis*; (*72*) for *Macaca assamensis*; (*73*) for *M. fuscata*; (*74*) for *Pongo pygmaeus wurmbii*; and *(75)* for *Varecia variegata*.

†Average frugivory based on (*70*) for *A. geoffroyi*; (*76*) for *C. ascanius* and *C. mitis*; (*77*) for *C. guereza*; (*78*) for *L. albigena* (reported as *Cercocebus albigena johnstoni*); (*79*) for *P. anubis*; (*80*) for *M. assamensis*; (*81*) for *M. fuscata*; (*82*) for *P. pygmaeus wurmbii*; and *(83)* for *V. variegata*.

‡Values based on mean ethanol content per fruit species.

§Values based on maximum ethanol content per fruit species.

Table S9. List of all model formulas used in statistical analyses. Each model is listed along with its type, distribution, and link function for the Phylogenetic, Environmental, and Canopy versus ground analyses. For each of the Environmental and Canopy versus ground analyses, listed models were compared with the corrected Akaike information criterion (AIC_c_) and the most parsimonious model for prediction selected. The term ‘scale(x)’ standardizes the predictor variable by converting it to *Z* scores, and the term ‘poly(x, 2)’ denotes a combination of linear and second order orthogonal polynomial predictors (where ‘x’ represents the dependent variable). The term ‘exp(pos + 0|1)’ applies random spatial terms to subsets of data with positive ethanol concentrations using a Euclidean distance matrix of sampling locations (based on latitude/longitude coordinates). The term ‘x1 * x2’ denotes an interaction term that includes the main effects of both ‘x1’ (independent variable 1) and ‘x2’ (independent variable 2), as well as their interaction. Random effects are specified as ‘(1 | x)’, where ‘x’ represents the grouping variable. Unique fruit ID and unique sample ID are included as random effects in the Canopy versus Ground models to account for unequal numbers of breathalyzer measurements among individual fruits and individual samples of fruit, respectively (which were not the same in cases where portions of the same fruit were sampled separately). Abbreviations: EtOH, ethanol; BR, biological realm; T, annual mean temperature; P, annual precipitation; PS, precipitation seasonality; NDVI, mean Normalized Difference Vegetation Index; PGLS, Phylogenetic Generalized Least Squares; MLE, Maximum Likelihood Estimation; GLMM, Generalized Linear Mixed Model.

| Analysis | Model number | Model formula | Model type | Model distribution (link function) |
| --- | --- | --- | --- | --- |
| Phylogenetic | 1 | Mean EtOH ~ 1 | PGLS with MLE of λ | Normal (identity) |
|  | 1 | Median EtOH ~ 1 | PGLS with MLE of λ | Normal (identity) |
|  | 1 | Maximum EtOH ~ 1 | PGLS with MLE of λ | Normal (identity) |
| Environmental (ethanol occurrence) | 1 | EtOH occurrence ~ -1 + BR + scale(T) + scale(P) + scale(PS) + scale(NDVI) + exp(pos + 0\|1) | Zero-inflated GLMM | Binomial (logit) |
|  | 2 | EtOH occurrence ~ -1 + BR + poly(T, 2) + poly(P, 2) + poly(PS, 2) + poly(NDVI, 2) + exp(pos + 0\|1) | Zero-inflated GLMM | Binomial (logit) |
|  | 3 | EtOH occurrence ~ 1 + scale(T) + scale(P) + scale(PS) + scale(NDVI) + exp(pos + 0\|1) | Zero-inflated GLMM | Binomial (logit) |
|  | 4* | EtOH occurrence ~ 1 + poly(T, 2) + poly(P, 2) + poly(PS, 2) + poly(NDVI, 2) + exp(pos + 0\|1) | Zero-inflated GLMM | Binomial (logit) |
|  | 5 | EtOH occurrence ~ 1 + scale(T) + scale(PS) + exp(pos + 0\|1) | Zero-inflated GLMM | Binomial (logit) |
|  | 6 | EtOH occurrence ~ 1 + BR + scale(T) + scale(P) + exp(pos + 0\|1) | Zero-inflated GLMM | Binomial (logit) |
|  | 7 | EtOH occurrence ~ -1 + BR + scale(T) + exp(pos + 0\|1) | Zero-inflated GLMM | Binomial (logit) |
|  | 8 | EtOH occurrence ~ poly(T, 2) + poly(P, 2) + poly(NDVI, 2) + exp(pos + 0\|1) | Zero-inflated GLMM | Binomial (logit) |
|  | 9 | EtOH occurrence ~ BR + poly(T, 2) + poly(P, 2) + poly(NDVI, 2) + exp(pos + 0\|1) | Zero-inflated GLMM | Binomial (logit) |
|  | 10 | EtOH occurrence ~ -1 + BR + poly(T, 2) + exp(pos + 0\|1) | Zero-inflated GLMM | Binomial (logit) |
|  | 11 | EtOH occurrence ~ -1 + BR + exp(pos + 0\|1) | Zero-inflated GLMM | Binomial (logit) |
|  | 12 | EtOH occurrence ~ poly(T, 2) + exp(pos + 0\|1) | Zero-inflated GLMM | Binomial (logit) |
| Environmental (mean ethanol concentration) | 1 | Mean EtOH ~ -1 + BR + scale(T) + scale(P) + scale(PS) + scale(NDVI) + exp(pos + 0\|1) | Zero-inflated GLMM | Gamma (log) |
|  | 2 | Mean EtOH ~ -1 + BR + poly(T, 2) + poly(P, 2) + poly(PS, 2) + poly(NDVI, 2) + exp(pos + 0\|1) | Zero-inflated GLMM | Gamma (log) |
|  | 3 | Mean EtOH ~ scale(T) + scale(P) + scale(PS) + scale(NDVI) + exp(pos + 0\|1) | Zero-inflated GLMM | Gamma (log) |
|  | 4 | Mean EtOH ~ poly(T, 2) + poly(P, 2) + poly(PS, 2) + poly(NDVI, 2) + exp(pos + 0\|1) | Zero-inflated GLMM | Gamma (log) |
|  | 5* | Mean EtOH ~ poly(T, 2) + poly(P, 2) + poly(PS, 2) + exp(pos + 0\|1) | Zero-inflated GLMM | Gamma (log) |
|  | 6 | Mean EtOH ~ -1 + BR + exp(pos + 0\|1) | Zero-inflated GLMM | Gamma (log) |
|  | 7 | Mean EtOH ~ T + exp(pos + 0\|1) | Zero-inflated GLMM | Gamma (log) |
|  | 8 | Mean EtOH ~ -1 + BR + poly(T, 2) + poly(P, 2) + exp(pos + 0\|1) | Zero-inflated GLMM | Gamma (log) |
|  | 9 | Mean EtOH ~ -1 + BR + poly(T, 2) + exp(pos + 0\|1) | Zero-inflated GLMM | Gamma (log) |
|  | 10 | Mean EtOH ~ poly(T, 2) + exp(pos + 0\|1) | Zero-inflated GLMM | Gamma (log) |
| Environmental (maximum ethanol concentration) | 1 | Maximum EtOH ~ -1 + BR + scale(T) + scale(P) + scale(PS) + scale(NDVI) + exp(pos + 0\|1) | Zero-inflated GLMM | Gamma (log) |
|  | 2 | Maximum EtOH ~ -1 + BR + poly(T, 2) + poly(P, 2) + poly(PS, 2) + poly(NDVI, 2) + exp(pos + 0\|1) | Zero-inflated GLMM | Gamma (log) |
|  | 3 | Maximum EtOH ~ scale(T) + scale(P) + scale(PS) + scale(NDVI) + exp(pos + 0\|1) | Zero-inflated GLMM | Gamma (log) |
|  | 4 | Maximum EtOH ~ poly(T, 2) + poly(P, 2) + poly(PS, 2) + poly(NDVI, 2) + exp(pos + 0\|1) | Zero-inflated GLMM | Gamma (log) |
|  | 5* | Maximum EtOH ~ poly(T, 2) + poly(P, 2) + poly(PS, 2) + exp(pos + 0\|1) | Zero-inflated GLMM | Gamma (log) |
|  | 6 | Maximum EtOH ~ -1 + BR + exp(pos + 0\|1) | Zero-inflated GLMM | Gamma (log) |
|  | 7 | Maximum EtOH ~ T + exp(pos + 0\|1) | Zero-inflated GLMM | Gamma (log) |
|  | 8 | Maximum EtOH ~ -1 + BR + poly(T, 2) + poly(P, 2) + exp(pos + 0\|1) | Zero-inflated GLMM | Gamma (log) |
|  | 9 | Maximum EtOH ~ -1 + BR + poly(T, 2) + exp(pos + 0\|1) | Zero-inflated GLMM | Gamma (log) |
|  | 10 | Maximum EtOH ~ poly(T, 2) + exp(pos + 0\|1) | Zero-inflated GLMM | Gamma (log) |
| Canopy versus ground | 1 | Proportion EtOH ~ Canopy location + (1 \| Fruit species name) + (1 \| Fruit intactness when measured) + (1 \| Unique sample ID) + (1 \| Unique fruit ID) | Zero-inflated GLMM | Beta (logit) |
|  | 2* | Proportion EtOH ~ Canopy location + (1 \| Fruit species name) + (1 \| Fruit intactness when measured) + (1 \| Unique sample ID) + (1 \| Unique fruit ID) | GLMM | Beta (logit) |
| Canopy versus ground (with ripeness stage) | 1 | Proportion EtOH ~ Canopy location * Ripeness stage + (1 \| Fruit species name) + (1 \| Fruit intactness when measured) + (1 \| Unique sample ID) + (1 \| Unique fruit ID) | Zero-inflated GLMM | Beta (logit) |
|  | 2* | Proportion EtOH ~ Canopy location * Ripeness stage + (1 \| Fruit species name) + (1 \| Fruit intactness when measured) + (1 \| Unique sample ID) + (1 \| Unique fruit ID) | GLMM | Beta (logit) |

__________________________

*Most supported model based on the AIC_c_.
